# A spatial multi-omics atlas of the human lung reveals a novel immune cell survival niche

**DOI:** 10.1101/2021.11.26.470108

**Authors:** Elo Madissoon, Amanda J. Oliver, Vitalii Kleshchevnikov, Anna Wilbrey-Clark, Krzysztof Polanski, Ana Ribeiro Orsi, Lira Mamanova, Liam Bolt, Nathan Richoz, Rasa Elmentaite, J. Patrick Pett, Ni Huang, Peng He, Monika Dabrowska, Sophie Pritchard, Liz Tuck, Elena Prigmore, Andrew Knights, Agnes Oszlanczi, Adam Hunter, Sara F. Vieira, Minal Patel, Nikitas Georgakopoulos, Krishnaa Mahbubani, Kourosh Saeb-Parsy, Menna Clatworthy, Omer Ali Bayraktar, Oliver Stegle, Natsuhiko Kumasaka, Sarah A. Teichmann, Kerstin B. Meyer

## Abstract

Multiple distinct cell types of the human lung and airways have been defined by single cell RNA sequencing (scRNAseq). Here we present a multi-omics spatial lung atlas to define novel cell types which we map back into the macro- and micro-anatomical tissue context to define functional tissue microenvironments. Firstly, we have generated single cell and nuclei RNA sequencing, VDJ-sequencing and Visium Spatial Transcriptomics data sets from 5 different locations of the human lung and airways. Secondly, we define additional cell types/states, as well as spatially map novel and known human airway cell types, such as adult lung chondrocytes, submucosal gland (SMG) duct cells, distinct pericyte and smooth muscle subtypes, immune-recruiting fibroblasts, peribronchial and perichondrial fibroblasts, peripheral nerve associated fibroblasts and Schwann cells. Finally, we define a survival niche for IgA-secreting plasma cells at the SMG, comprising the newly defined epithelial SMG-Duct cells, and B and T lineage immune cells. Using our transcriptomic data for cell-cell interaction analysis, we propose a signalling circuit that establishes and supports this niche. Overall, we provide a transcriptional and spatial lung atlas with multiple novel cell types that allows for the study of specific tissue microenvironments such as the newly defined gland-associated lymphoid niche (GALN).

## Introduction

Lung disease is the third-largest cause of mortality worldwide (Angelidis et al. 2019)), making a better understanding of the cell types that define lung function imperative. Histological definition of the lungs identified around 45 different cell types (Gehr, Bachofen, and Weibel 1978; Ross and Pawlina 2006), ranging from well-characterised lung epithelial cells to rare neuroendocrine and tuft cells and varying by location from the trachea to the small airways and alveoli (Boers, Ambergen, and Thunnissen 1998; Travaglini et al. 2020). In addition to its respiratory function, the lung has an important barrier function. Whilst other mucosal barrier tissues such as the gut and nasopharynx orchestrate adaptive immunity through well defined mucosa-associated lymphoid tissue (MALT), such secondary lymphoid structures have not been reported in the healthy human lung (Kato et al. 2013).

The LungMAP and Human Lung Cell Atlas consortia (Schiller et al. 2019; Ardini-Poleske et al. 2017), have harnessed recent technology advances in single cell and single nucleus RNA sequencing (Wilbrey-Clark, Roberts, and Teichmann 2020) and generated a number of atlases characterising lung cell types in mice (Han et al. 2018; Tabula Muris Consortium et al. 2018; Montoro et al. 2018) and humans (Travaglini et al. 2020; Angelidis et al. 2019; Plasschaert et al. 2018; Vieira Braga et al. 2019; Madissoon et al. 2019). In addition to previously-known classifications, this work has identified new cell states and types such as the pulmonary ionocyte (Plasschaert et al. 2018; Montoro et al. 2018), deuterosomal cells (Ruiz García et al. 2019) and a tissue migratory T helper cell population (Vieira Braga et al. 2019). Respiratory system diseases such as asthma (Vieira Braga et al. 2019), pulmonary fibrosis (Habermann et al. 2020; Reyfman et al. 2019; Adams et al. 2020), chronic rhinosinusitis (Ordovas-Montanes et al. 2018), cystic fibrosis (Carraro et al. 2021) and the effect of smoking (Ordovas-Montanes et al. 2018; Goldfarbmuren et al. 2020a), have all been studied at the single cell transcriptome level.

However, unbiased spatial mapping of cell types back into the tissue context is still largely lacking, and enzymatic dissociation procedures can lead to unrepresentative cell type retrieval. To overcome these limitations, we combined single cell and single nuclei RNA sequencing with Visium Spatial Transcriptomics (ST) to create a lung cell atlas of 129,340 cells, 63,768 nuclei and 16 tissue sections from human trachea, bronchi and parenchyma. We describe the transcriptional signatures of 77 cell types and states, including for the first time that of human airway chondrocytes, two types of Schwann cells, nerve-associated cells, submucosal gland (SMG) duct cells, and multiple perivascular, fibroblast and macrophage subsets. Our deep tissue profiling coupled with spatial gene expression profiling has allowed us to identify a new immune niche for IgA-secreting plasma cells at the SMG of the airways. We predict cell-cell interactions across endothelial, epithelial, stromal and immune cells that help to establish this niche. IgA secretion plays a critical role in preventing respiratory infections, and hence our findings are relevant for a deeper understanding of a wide range of respiratory infectious diseases, including COVID-19 (Sterlin et al. 2021).

## Results and Discussion

### A multi-omic transcriptional atlas of the human lung and airway with spatial resolution

To generate a comprehensive representation of the cell types in human lung and airway we applied single cell and nuclei RNAseq (scRNAseq, snRNAseq), VDJ sequencing and spatial transcriptomics to deep tissue samples (Figure 1a, Tables S1-3). Tissue pieces (1-2 cm^3^) from healthy donors were collected from the trachea, bronchi at the 1st or 2nd generation airway (Bronchi 2-3), bronchi at the 3rd to 4th generation airway (Bronchi 4), parenchyma from the upper left lobe (UpperPar) and the lower left lobe (LowerPar) to capture deep structures such as cartilage, muscle and the submucosal glands in the airways (Figure S1a).

**Figure 1.**
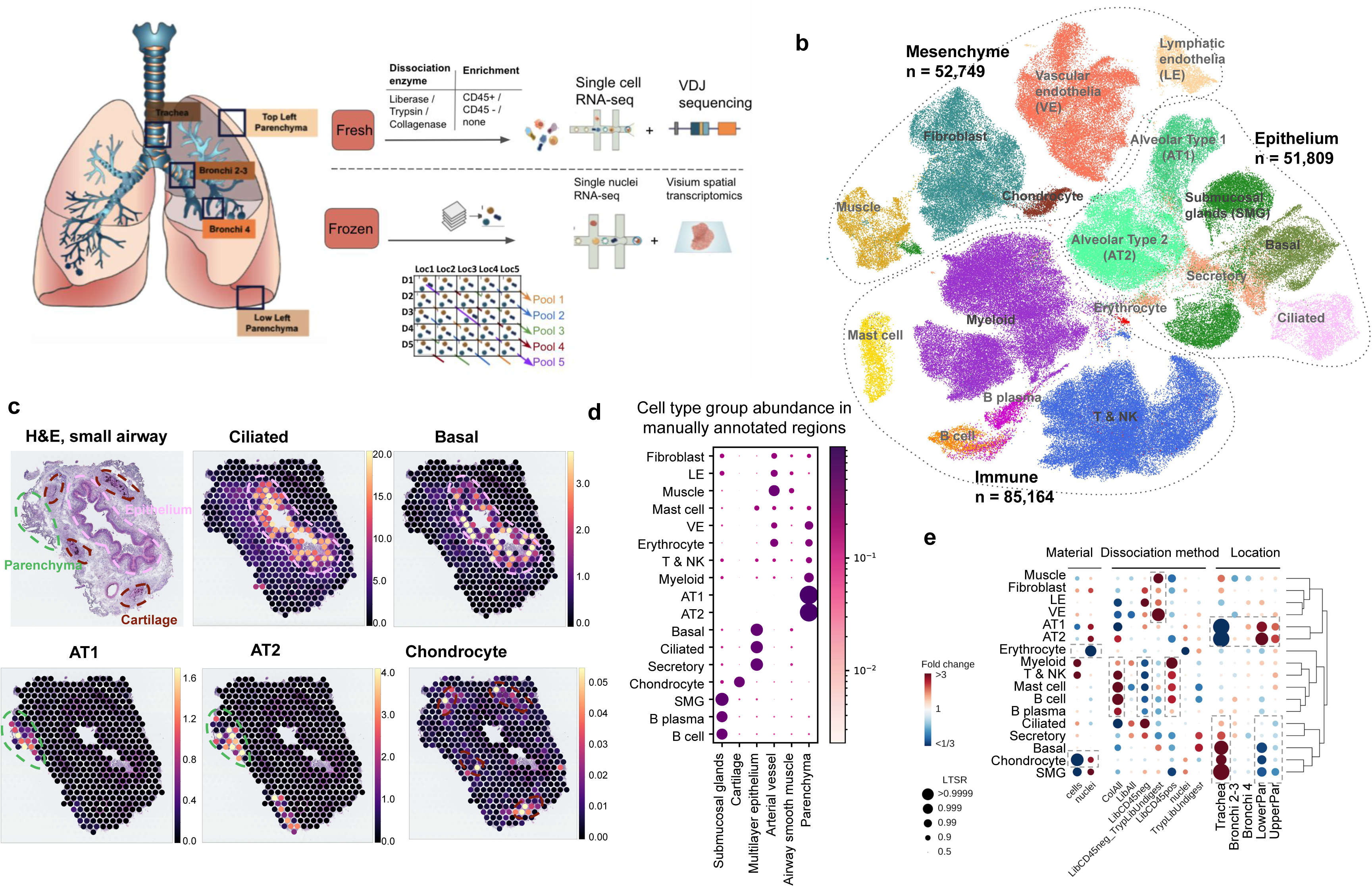
Spatial multi-omics atlas of the human lung allows the identification of cell types and their location. (a) Multi-omics spatial lung atlas experimental design included fresh and frozen sampling from 5 locations for single-cell (sc) RNA sequencing (RNAseq), sc VDJ-sequencing, single-nuclei (sn) RNAseq and Visium Spatial Transcriptomics (ST). Five donors (D) from the frozen samples were pooled into five reactions, each containing different locations (Loc) from donors. (b) sc/sn RNA-seq captures cell types from all cell type groups. (c) Cell2location mapping on Visium ST from small airway shows matching of cell types to expected structures. H&E staining of small airway section and cell abundance estimated by cell2location for ciliated, basal epithelium, AT1, AT2 and chondrocyte cell types with histology image in the background. Dotted lines circle the epithelium (pink), parenchyma (green) and cartilage (brown). (d) Cell type groups are enriched in expected regions on Visium ST across sections. Normalised average cell abundance (dot size and color) for cell types annotated in (b), across the manually annotated regions in the Visium data. (e) Cell types capture is affected by protocol and location. Cell type proportion analysis with fold changes and Local True Sign Rate (LTRS) score for all cell type groups with regards to the material, protocol and location. Dashed boxes highlight the greatest changes. AT1 - alveolar type 1, AT2 - alveolar type 2.

In total 193,108 good quality transcriptomes were annotated into transcriptionally distinct broad cell type groups: epithelial (secreting, basal, alveolar type 1 and 2 - AT1 & AT2 - pneumocytes, ciliated, and SMG cells), immune (lymphoid, myeloid, B-cells, plasma, mast cells), erythrocytes, endothelial (vascular- VE, lymphatic - LE) and stromal cells (smooth muscle, fibroblasts, mesothelial and chondrocytes) according to widely accepted marker genes (Figure 1b, Figure S1b). We combined the analysis of suspension cells and nuclei with Visium Spatial Transcriptomics (ST) analysis to map the identified cell types back to tissue structures on 16 tissue sections from 5 locations (Figure S2) which were manually annotated into 13 regions (Table S4). As expected, we mapped ciliated epithelial cells to the lumen of the airway surrounded by basal cells, and AT1 and AT2 to lung parenchyma (Figure 1c, d), using the Cell2location (Kleshchevnikov et al., n.d.) algorithm. In order to examinethe benefits of different protocols and sampling locations, we used a Poisson linear mixed model to assess the relative effects of these variables on cell type composition (see Methods) (Figure 1e). As expected, different enzymatic treatments enriched for specific cell type groups but had little effect, less than 1% of the total variance, on gene expression (Figure S1 h).

As a demonstration of the depth and novelty of our atlas, we transcriptionally defined chondrocytes for the first time in human lungs (*ACAN, CHAD, COL9A3, HAPLN1* and *CYTL1)* (Figure S1 c, d) and mapped these cells to the cartilage with Visium ST (Figure 1 c, d). Chondrocytes were mostly released using single nuclei sequencing from trachea (Figure 1 e, Figure S1 e-g), demonstrating the utility of our multi-omics, multi-location atlasing of the human lung.

Overall, we generated a large scRNAseq and snRNAseq dataset with wide representation of distinct cell lineages and matching spatial gene expression data, allowing further exploration of cellular heterogeneity and possible cell-cell interactions to better understand lung function.

#### Identification of rare fibroblasts with immune recruiting properties

The sequential clustering of lung and airway fibroblasts identified 11 cell types with distinct marker genes (Figure 2a, Figure S3 a, b). We annotate previously described myofibroblasts, mesothelial, adventitial and alveolar fibroblasts (Travaglini et al. 2020; Xia et al. 2014; Gomez et al. 2016; Bauer et al. 2015; Kanamori-Katayama et al. 2011), as well as multiple novel and less studied cell types that were not predicted with existing annotations (Figure S3 c, d). While most of the cell types were isolated from all locations and protocols, there were clear differences in cell type proportions (Figure 2b, Figure S3b). As expected, the adventitial fibroblasts (Fibro-adv) were enriched in the airway samples, while the alveolar fibroblasts (Fibro-alv) were depleted. We further observed a rare cell type recovered only from the single cells, but not from single nuclei. This rare type expressed the chemokines CCL19 and CCL21, also highly expressed in fibroblast reticular cells responsible for T-cell positioning in the lymph nodes, and GREM1 (Kapoor et al. 2021), CXCL13 and FDCSP, a follicular dendritic cell (FDC) marker responsible for B-cell positioning (Wang et al. 2011; Marshall et al. 2002)(Figure 2c). These cells mapped to an area of immune cell infiltration on the bronchus with Visium ST, as well as with smFISH methods (Figure 2d, Figure S4 a, b) and were therefore annotated as immune recruiting fibroblasts (IR-fibro). Immune infiltrates were only found in a proportion of the donors and sections, but in keeping with the expected frequency in donors with and without smoking history (Elliot et al. 2004). IR-fibros display some characteristics of fibroblasts from secondary and tertiary lymphoid structures, and this similarity is further supported by our finding that the gene signature of germinal center fibroblast cell types from the Peyer’s Patches of human gut (R. Elmentaite et al. 2021) also map to the area of the immune infiltrate on lung Visium ST sections (Figure S4b). In conclusion we describe an IR-fibro population with a likely role in immune cell recruitment in the healthy bronchus.

**Figure 2.**
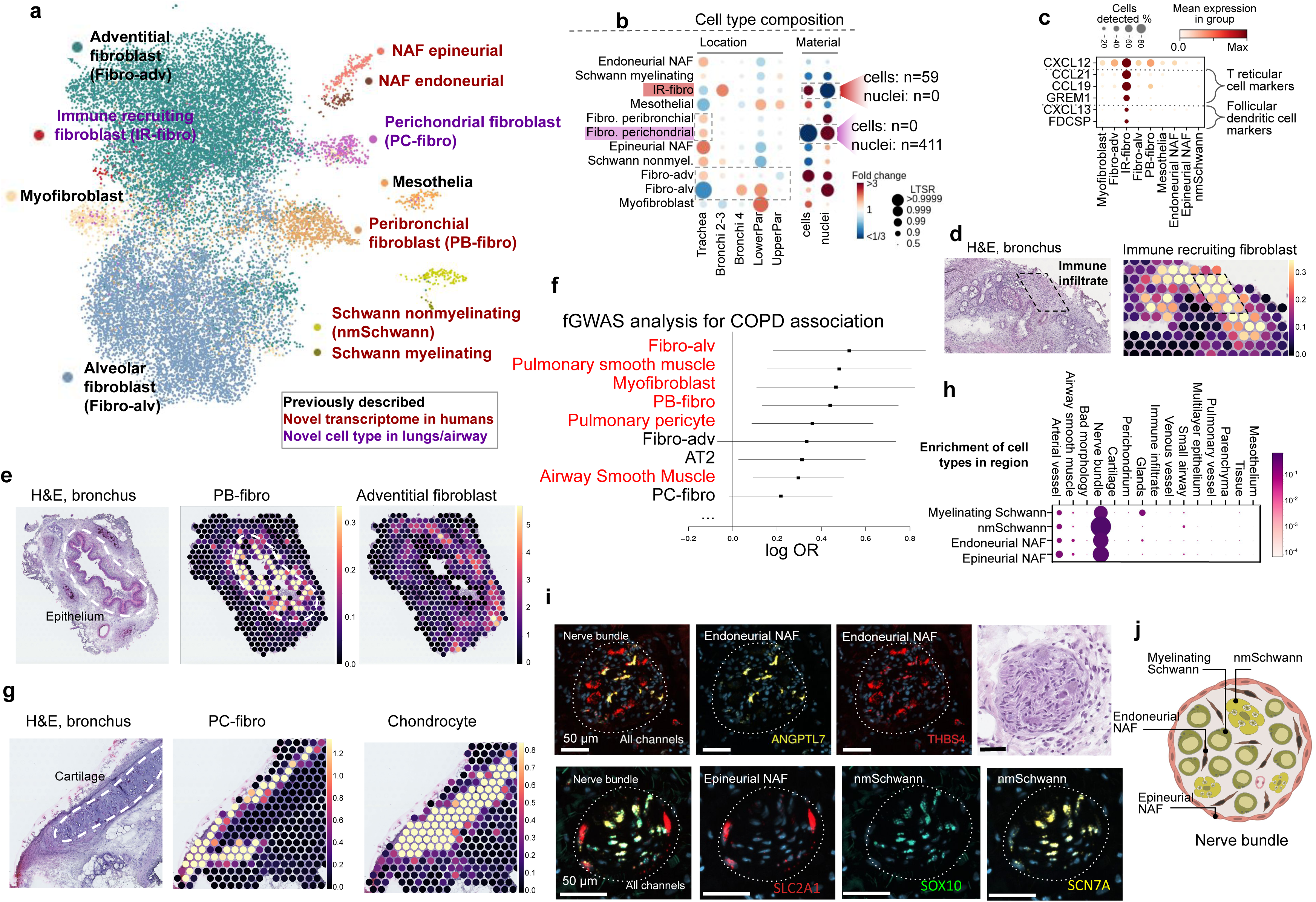
High resolution of lung and airway fibroblasts and their spatial location. (a) Sequential clustering reveals 11 fibroblast populations in airways and lungs on UMAP. (b) Sample collection location and processing method affect recovered cell type proportions. Cell type composition analysis with fibroblast compartment, including location, material, dissociation protocol and donor in the model. (c) Markers of immune recruiting fibroblast population (IR-fibro) overlap with T reticular and Follicular Dendritic cell markers. (d) Immune recruiting fibroblast co-localise with immune infiltrate region in the airways with Cell2location anaysis. (e) Peribronchial fibroblasts (PB-fibro) co-localise around small airway. (f) Peribronchial fibroblasts are associated with COPD with fGWAS analysis. (g) Perichondrial fibroblasts (PC-fibro) localise around the cartilage regions in the airway. (h) Scwhann cells and nerve-associated fibroblasts (NAF) co-localise with peripheral nerve bundles via smFISH method. (i) Nerve-associated cell types markers have distinct locations in the airway nerve bundles. (j) Schematic representation of the described nerve-associated populations in the peripheral nerves of the airway.

#### Perichondrial and peribronchial fibroblasts reveal disease associations

We next examined two fibroblast populations enriched in trachea: peribronchial (PB-fibro) and perichondrial (PC-fibro) fibroblasts (Figure 2b). For the first time we provide the human transcriptome for PB-fibro with COL15A1 and ENTPD1 markers and localisation around the airway epithelium (Figure 2e, Figure S5a). Functional GWAS analysis, that quantifies systematic associations between cell-type specific gene expression and disease-associated SNPs (R. Elmentaite et al. 2021), demonstrated that PB-fibros are linked to chronic obstructive pulmonary disease (COPD). PB-fibro also express LGR5, as reported for mouse (Tsukui et al. 2020; J.-H. Lee et al. 2017), which suggests potential for analogous mesenchymal stem cell function in humans (Figure S5b). A COPD-related increase in WNT-5A downregulates Lgr5 (Baarsma et al. 2017), which suggests PB-fibro could be targeted therapeutically to inhibit regenerative failure in COPD specifically in the airways.

The second airway-enriched cell type, PC-fibro, was detected only in the single nuclei data (Figure 2b), likely due to difficulty in dissociation of cartilage. The cell type was located around the chondrocytes within cartilage in our Visium ST (Figure 2g) and could be identified by the expression of the marker gene *COL12A1* in the human protein atlas (Figure S5c). Among the marker genes for PC-fibro, we detected bone development genes *LGR4* and *LGR6* (Doherty and Sanjay 2020) (Figure S5b). PC-fibro, which express standard fibroblasts markers, is unique in expressing genes relevant for bone development, placing these cells into a trajectory from adventitial fibroblasts to chondrocytes (Figure S5 d, e). Finally, PC-fibro markers were enriched in genes causing skeletal abnormalities in humans (Figure S5f), including FLNB and FGFR2 that are mutated in rare diseases causing skeletal malformations (Bochukova et al. 2009; Robertson 2008). In conclusion, we have identified perichondrial fibroblasts around the healthy human airway cartilage.

#### Resolution of four distinct cell types in the peripheral nerves from the airway

Finally, among the fibroblast compartment we identify four rare cell clusters within the peripheral nerves in the airway: myelinating Schwann cells (*NFASC*, *NCMAP*, *MBP*, *PRX)*, non-myelinating Schwann cells (nmScwann) *(NGFR*, *SCN7A*, *CHD2*, *L1CAM*, *NCAM1)* (Chen et al. 2021; Gerber et al. 2021; Renthal et al. 2020; Wolbert et al. 2020), endoneurial nerve-associated fibroblasts (NAF) (SOX9, OSR2) (Chen et al. 2021) and epineurial NAF (Figure S6a). Localisation of those cell types in peripheral nerves is shown by bulk RNA-seq across tissues (Figure S6d), Visum ST (Figure 2h, Figure S6 e), protein staining (Figure S6 f-h) and smFISH (Figure 2i, Figure S6 i). Specifically, we show the expression of epineurial NAF around, and endoneurial NAF with Schwann cells inside the nerve bundle in human airway samples. This further confirms the identity of myelinating and non-myelinating Schwann cells with marker gene set enrichment in myelination and cell adhesion categories respectively (Figure S6 b, c). This analysis further reveals EVX1 - a homeobox gene in spinal cord development - as a potential regulator of myelinating Schwann cells. Second, the nmSchwann expressed some genes involved in Schwann cell development and maintenance (*SOX10*) and myelination (*ERBB3* and *LGI4*) (Figure S6 a, Figure 2i, Figure S6 i). Finally, we observed specific expression of rare peripheral nervous system (PNS) disease genes in these populations (Figure S6 j). Thus, our atlas discerns microanatomical location within structures such as peripheral nerves (Figure 2j) and pinpoints disease associations, even for very rare cell types. The cell types described here exist in other tissues; therefore this data could have implications outside the lung.

### Resolution of the vascular cell types in the systemic and pulmonary circulation

We next focused on studying of the vasculature, which in the lung consists of both systemic circulation providing oxygen to the tissue, and pulmonary circulation where gas exchange occurs. Uniquely in our dataset, we could distinguish clusters of pulmonary and systemic circulation by the distribution of cell types, where pulmonary cells are enriched in parenchyma and systemic cells are enriched in the trachea (Figure 3d, Supplementary Figure S7e). We observed previously described venous (E-Ven-syst, E-Ven-pulm) and capillary (Cap, Car4-Aerocyte) endothelial cell types (Figure 3a, Supplementary Figure S7a) (Vila Ellis et al. 2020; Niethamer et al. 2020; Gillich et al. 2020; Schupp et al. 2021). In addition, we now distinguish arterial endothelial cell types (systemic E-Art-syst and pulmonary E-Art-pulm), which have distinct localisation on Visium ST with E-Art-syst enriched in the trachea (Figure 3a, b). The smooth muscle cells included non-vascular airway smooth muscle cells (ASM), pulmonary perivascular cell types (smooth muscle SM-pulm and pericytes Peri-pulm) (Travaglini et al. 2020) and systemic perivascular cells (smooth muscle SM-Art-syst, pericytes Peri-syst and venous perivascular cells IR-Ven-Peri) (Figure 3a, c Supplementary Figure S7b). The capture of ASM was mainly enabled by single nuclei data (Figure 3d) that allowed for the detection of reliable marker genes for non-vascular smooth muscle cell types across tissues (Supplementary Figure S7 c, d).

**Figure 3.**
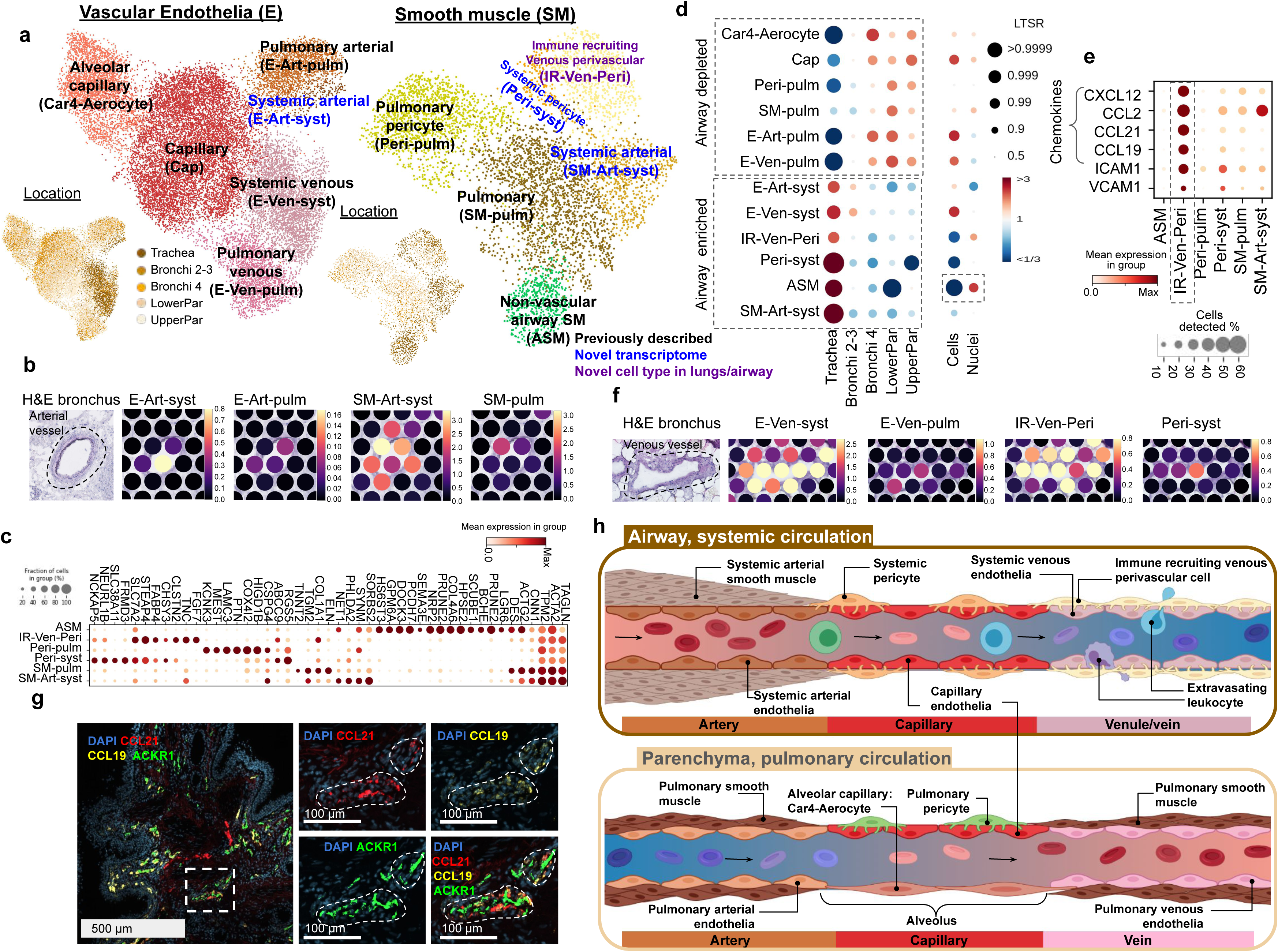
Cell types of the systemic and pulmonary circulation. (a) UMAP visualisation of scRNA-seq data from the vascular endothelia and smooth muscle compartments. Color of dots shows cell type or location. Color of text reflects novelty. (b) E-Art-syst co-localises with arterial vessel and SM-Art-syst in the airway. (c) Marker genes of the smooth muscle compartment. (d) Enrichment of cell types shows preferential localisation to the airways versus parenchyma and cells versus nuclei. (e) IR-Ven-peri expresses immune recruiting chemokines and cell-cell adhesion molecules. (f) E-Ven-sys and IR-Ven-Peri co-localise at a venous vessel in the airway. (g) IR-Ven-Peri markers localise adjacent to the venous vessel marker in the airway. (h) Resolution of transcriptionally defined vascular cells in the systemic and pulmonary circulations.

In addition, we identified another perivascular cell type in the airways, expressing ABCC and ICAM1, but not CSPG4 - similarly to the postcapillary venous perivascular cells with a role in the homing of immune cells in peripheral lymph nodes (Murfee, Skalak, and Peirce 2005; Proebstl et al. 2012) (Figure 3c). This cell type expressed known leukocyte-recruiting chemokines (Figure 3e) and co-localised with a venous vessel in the Visium ST bronchi section (Figure 3f) and in smFISH microscopy (Figure 3g, Supplementary Figure S7 f). These cells were therefore annotated as immune recruiting venous perivascular cells (IR-Ven-Peri). The potential lymphocyte recruiting capacity in both venous endothelial and IR-Ven-Peri (Supplementary Figure S7 g) suggests a role in lymphocyte extravasation in the airway veins, similar to the postcapillary venules in peripheral lymph nodes.

In summary, we distinguish between cells of the systemic and pulmonary circulation, and describe a novel immune recruiting postcapillary venous pericyte (IR-Ven-Peri). We provide marker genes for all of these populations, map these cells back into the tissue context and thereby further define the relationship between the endothelial and perivascular cells in the systemic and pulmonary circulation (Figure 3h).

#### Identification of duct and myoepithelial cells of the human airway submucosal glands (SMG)

Further analysis of the epithelial compartment (excluding AT1 and AT2 cells) identified expected ciliated, goblet, basal, suprabasal, dividing basal, club, SMG mucous and serous populations along with rare deuterosomal, ionocyte & brush and neuroendocrine cells. In addition to these, we identified a novel population between serous, mucous and club/goblet cells (Figure 4a, Figure S8 a, b) with marker genes overlapping with SMG duct cells in human oesophagus and preferential localisation in trachea (Madissoon et al. 2019; Nowicki-Osuch et al. 2021). Thus, we hypothesise that these cells are the columnar non-ciliated duct cells of the human airway, previously only characterised in mice at the single cell level (Widdicombe and Wine 2015; Meyrick, Sturgess, and Reid 1969; A. E. Hegab et al. 2012). smFISH staining for *ALDH1A3, MIA* and *RARRES1* validated the localisation at the SMG and distinguished these cells from the serous, mucous and other epithelial cells (Figure 4b, Figure S8 e, f). Cell2location integration of single-cell and Visium data could also distinguish distinct locations of SMG duct cells compared to SMG mucous and serous cells, providing orthogonal evidence of the identification of a new, distinct cell type (Figure S8h). In addition, Velocyto analysis suggested these cells lie on a trajectory towards the surface epithelial populations (Figure S8c), supporting a potential role in airway surface regeneration, as has been demonstrated in mice (Tata et al. 2018; Ahmed E. Hegab et al. 2011).

**Figure 4.**
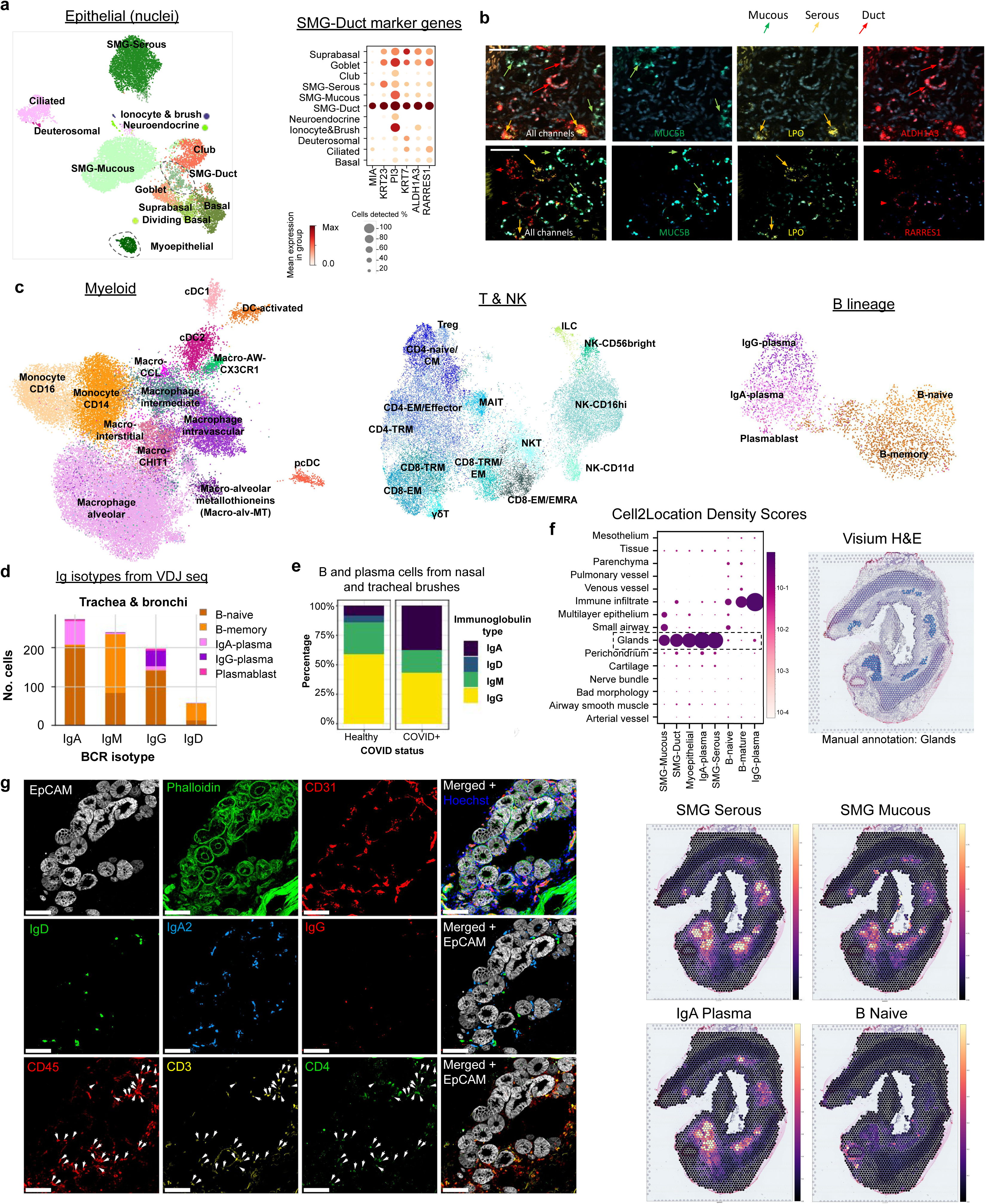
IgA plasma cells in human airways co-localise with submucosal glands. (a) UMAP of single nuclei RNA-seq airway epithelial cells (excluding parenchyma AT1 and AT2 epithelial cells) with dashed line around SMG-Duct and myoepithelial cells and a dot plot for the SMG-Duct marker genes. (b) RNAscope staining of mucous, serous and duct cells with respective marker genes MUC5B, LPO and ALDH1A3/RRARES1 in human bronchus. (c) UMAP plots of myeloid, T/NK and B cell lineage cells, colored by cell type. (d) Number of B lineage cells with different Ig isotypes in airway (trachea & bronchi) from the analysis of VDJ amplified libraries. (e) Percentages of different isotypes of B and plasma cells from the nasal and tracheal brushes of COVID-19+ and healthy control patients (Yoshida et al. 2021). Patients with over 20 B and plasma cells were considered. (f) Visium ST results show IgA plasma cells specific localisation in the glands. H&E on bronchi with manually annotated gland regions shown in blue, Cell2location density scores for IgA-plasma cells, and normalised average cell abundance (dot size and color) for SMG cell types and B lineage cell types across the manually annotated regions in the Visium data. (g) Multiplex IHC staining of human trachea for the SMG structure (Hoechst for nuclei, EpCAM for epithelium, Phalloidin for actin, CD31 for vessels), B lineage markers (IgD, IgA2, IgG) and CD4 T-cells (CD45, CD3, CD4). Arrows point to CD45+CD3+CD4+ cells. Scale bar 100 µm.

An additional population was revealed from snRNAseq, exhibiting both basal epithelium and muscle markers with localisation around the glands (Figure 4a, Figure S8g), consistent with a myoepithelial cell signature (Goldfarbmuren et al. 2020b). Co-localisation of *FHOD3*, epithelial marker *KRT14* and muscle marker *TAGLN* in the same cells confirmed the myoepithelial cell signature by smFISH (Figure S8g). Interestingly, mouse myoepithelial cells have also been shown to regenerate the surface airway epithelium (Tata et al. 2018)). This cell type is not as well described in humans, potentially due to difficulties in dissociating this cell type from the airways consistent with exclusive recovery of myoepithelial cells from nuclei data. Overall, we provide the human transcriptome for novel SMG populations, and mapped the constituents of the SMG and their spatial location using both spatial transcriptomics and smFISH.

#### Proportions and gene signatures of the airway epithelial cells

Using statistical modelling that accounted for material, donor and dissociation protocol (see Methods), we examined the proportions of airway epithelial cells at the 5 different locations sampled (Figure S8c), with significance given as local true sign rate (LTR). As expected, club cells were enriched in the parenchyma, whereas SMG epithelial cells were enriched in the trachea. While donor and protocol had little effect on the epithelial cell type proportions, the starting material of cells versus nuclei strongly influenced cell type composition (Figure S8c). SMG epithelial cell capture was increased in nuclei, while the capture of other epithelial cell types was better achieved with the dissociation into single cells (Figure S8c).

Taking advantage of our multi-location data, we compared gene expression for ciliated cells, as one of the most abundant cell types present, across the five locations. We avoided any artefacts due to differing ambient RNA contamination between locations using only snRNAseq data where samples were pooled across locations for the 10X sequencing reaction (Figure 1a). Using a linear mixed model (A. M. H. Young et al. 2021) (Methods) we detected 80 differentially expressed genes in ciliated cells from trachea compared to other locations (LTSR>0.9), many of which are upregulated in nasopharyngeal carcinoma including *FBXL7*, *TSHZ2* and *RAET1E* (Figure S8i) (Borchers et al. 2009; Motz et al. 2010; Wortham et al. 2013). We also examined *ACE2* expression, a SARS-COV-2 entry gene, which we found to be highest in ciliated cells from the trachea, and lower in the distal regions of the lung, where expression of *ACE2* is likely to be more relevant in AT2 cells as reported previously (Figure S8j) (Deprez et al. 2020).

Altogether, we describe transcriptomes for two key cell types involved in the SMG structure in the human airways which also functions as an immunological niche which we describe below.

### Immune cells in the lung and airways

#### Myeloid cells show previously undescribed heterogeneity

For the immune compartment, we identified all major populations including myeloid, T&NK, B lineage, mast cells and megakaryocytes which were analysed separately to reveal previously undescribed heterogeneity, especially in the myeloid cells (Figure 4c, Figure S9a-c, Figure S10 a). We found all major myeloid cell types (DCs, monocytes and macrophages) including many known and novel subsets. Previously identified macrophage subsets included intravascular macrophages (expressing *LYVE1* and *MAF*) (Chakarov et al. 2019; Evren et al. 2021), Macro-AW-CX3CR1 (Chakarov et al. 2019; Pirzgalska et al. 2017; Y. Wolf et al. 2017; Hulsmans et al. 2017), Macro-CHIT1 (CHIT1 expressing with roles in asthma, COPD and lung fibrosis) (Travaglini et al. 2020; Chang, Sharma, and Dela Cruz 2020) and interstitial macrophages (Macro-interstitial expressing chemokines *CXCL9, 10* and *11*)(Evren et al. 2021). We identified a new cluster expressing both monocyte CD14 and macrophage markers, termed Macro-intermediate (Figure S9a). Among alveolar macrophages, two more clusters appeared: dividing cells (Macro-alv-dividing), and a novel cell cluster expressing metallothioneins (Macro-alv-MT) including *MT1G*, *MT1X* and *MT1F*. Metallothioneins have a role in binding and metabolising metal ions (Artur Krężel 2017), immunity and stress response (Subramanian Vignesh and Deepe 2017; Takano et al. 2004), and therefore this population may have a function in response to air pollution. The final rare novel population of macrophages expressed chemokines including *CXCL8*, *CCL4* and *CCL20* and was named Macro-CCL. While the expression of *CCL4* was previously identified in interstitial macrophages (Evren et al. 2021), the expression of *CXCL8* and *CCL20* distinguishes this novel subset. Dysregulation of *CXCL8* expression is associated with multiple lung conditions including infection, asthma, IPF and COPD (Mukaida 2003) and was identified as a marker for a separate macrophage population in psoriatic skin (Reynolds et al. 2021).

Overall, we have identified multiple known and novel myeloid populations in the healthy human lungs and airways, many of them expressing specific sets of chemokines, orchestrating the complex lung immune homeostasis.

#### Different subsets of T & NK cells in the lung and airways

T lymphocytes and natural killer (NK) cells included all major cell types (CD4, CD8, mucosal-associated invariant T (MAIT), NK, NKT, innate lymphoid cells (ILC)) and their subsets (Figure 4c, Figure S9b). In the CD4 compartment we distinguished naive/central memory (CD4-naive/CM), effector memory/effector (CD4-EM/Effector), regulatory T cells (Treg) and tissue resident memory (CD4-TRM) cells. Within CD8 cells we found gamma-delta T cells (γδT) TRMs (CD8-TRM) (Hadley et al. 1997) which nicely localised to airway epithelium in our spatial data as known in the literature (Figure S9d)(Piet et al. 2011; Wu et al. 2014). In addition we saw two distinct CD8+ clusters analogous to populations found in the lung in cross tissue analysis (Conde et al., n.d.) expressing *CX3CR1* and *GZMB* (CD8-EM/EMRA) and *CRTAM* and *GZMK* (CD8-TRM/EM). NK subsets included clusters with markers *ITGAD*/CD11d, *LAG3*, *KLRC3*/NKG3E, *KLRC2*/NKG2C (NK-CD11d), CD16+ (NK-CD16hi) and CD56 bright NK cells (NK-CD56 bright) (Ghilas et al., n.d.; Böttcher et al. 2018). NK cells positive for CD11d have an activation or viral response in infection in both mouse and human (Ma et al. 2017; Fang et al. 2011; Triebel et al. 1990) and were previously shown in human blood (Siegers et al. 2017). To our knowledge this is the first time this subset has been described in healthy human lung.

The T and NK cells displayed striking donor-to-donor variability in cell type proportions compared to the myeloid clusters (Figure S9 g-i), consistent with higher inter-individual variability in the adaptive immune compartment. The location of origin, material and protocol explained little variation for any of the cell types proportions, including for the B cells.

We also obtained TCR VDJ sequencing data that confirmed MAIT cell type annotation (with preferential use of TRAJ33 and TRAV1-2)(Treiner et al. 2003), and showed low clonal expansion in naive and Treg populations compared to memory and effector subsets (Figure S9 e). Lastly we show that, as expected, there was no clonal sharing between individuals, but expanded clones were found in multiple locations of the lung within one donor (Figure S9 f).

#### Co-localisation of IgA plasma cells with the SMG

B cells included naive and memory B cells, and plasma cells that were further annotated into immunoglobulin (Ig) IgG or IgA secreting plasma cells and plasmablasts (Figure 4c, Figure S10a, b), and this annotation was supported by VDJ-seq data via Scirpy BCR isotype analysis. IgA, which is important for mucosal immunity (Corthesy 2013, Kunkel and Butcher 2003, Salvi and Holgate 1999), was the most frequent isotype in the airway samples, while only the third most abundant in the parenchyma (Figure 4d, Figure S10 e). Interestingly, we observe that proportions of IgA plasma cells, relative to IgG, are increased in COVID-19 patients versus healthy controls in single cell data from nasal and bronchial brush samples (Figure 4e) (Yoshida et al. 2021). The distinguishing markers for IgA versus IgG plasma cells included *CCR10* and B-cell maturation antigen BCMA (*TNFRSF17*) (Figure S10 b), which are important for plasma cell localisation and survival, respectively (Morteau et al. 2008; Kunkel and Butcher 2003; O’Connor et al. 2004)

Using Visium ST, we observed the localisation of IgA B plasma cells, but not B or IgG plasma cells, in the airway SMG in trachea and bronchi sections (Figure 4f). Annotation of anatomical regions across all Visium sections confirmed co-localisation of IgA plasma cells with duct, serous and mucous SMG cells, whilst IgG mapped to immune infiltrates (Figure 4f). The mapping of IgA plasma cells to the SMG is confirmed in the Human Protein Atlas which shows an abundance of plasma cells (MZB1+) in the SMG region of the bronchus and the nasopharynx (Figure S10 c, d), along with a study from the 1970s that showed localisation of IgA plasma cells in human airway SMG (Soutar 1976).

Using multiplex IHC we showed the specific presence of IgA2 plasma cells in the SMG at single cell resolution, while IgG positive cells were present in the airways only outside the SMG, consistent with Visium ST (Figure 4g, Figure S10 f). We also detected IgD+ naive B cells and CD3+CD4+ T helper cells in the SMG (Figure 4g), suggesting that IgA plasma cells are supported by a complement of cell types that can orchestrate B cell maturation for IgA secretion directly into the airway mucous. We hypothesise that together these different cell types constitute an immune niche which we term gland associated lymphoid niche (GALN). Understanding the immunological mechanisms at the SMG can help understand disease, as increased plasma cell numbers in SMG have been shown in smokers (Soutar 1976), in COPD (Zhu et al. 2007) and in Kawasaki disease (Rowley et al. 2000).

#### Cell-cell interactions and the SMG immune cell niche

To understand colocalization of B cells, IgA plasma cells and T cells in the SMG (Figure 4f), we explored the molecular mechanisms underpinning the SMG as a potential immune niche. We report that expression of pIgR, which facilitates transcytosis of polymeric Ig across the surface epithelium, was high across all SMG epithelial cells, as was Mucosal Epithelial Chemokine (MEC)/CCL28, known to recruit IgA plasma cells through CCR10 in other mucosal sites (Figure 5a, b) (Wilson and Butcher 2004; Morteau et al. 2008). Using cell-cell interaction analysis tool CellChat on cells from the airways (Jin et al. 2021) we saw that, in addition to the CCL28-CCR10 axis between SMG cells and B plasma cells (combined IgA, IgG and plasmablasts), SMG-Duct cells were predicted to interact with CCR6 on memory and naive B cells and CD4 T cells (combined CD4 subsets, excluding Tregs) through CCL20 (A. Y. S. Lee et al. 2017; Elgueta et al. 2015; Bowman et al. 2000) (Figure 5b-d).

**Figure 5.**
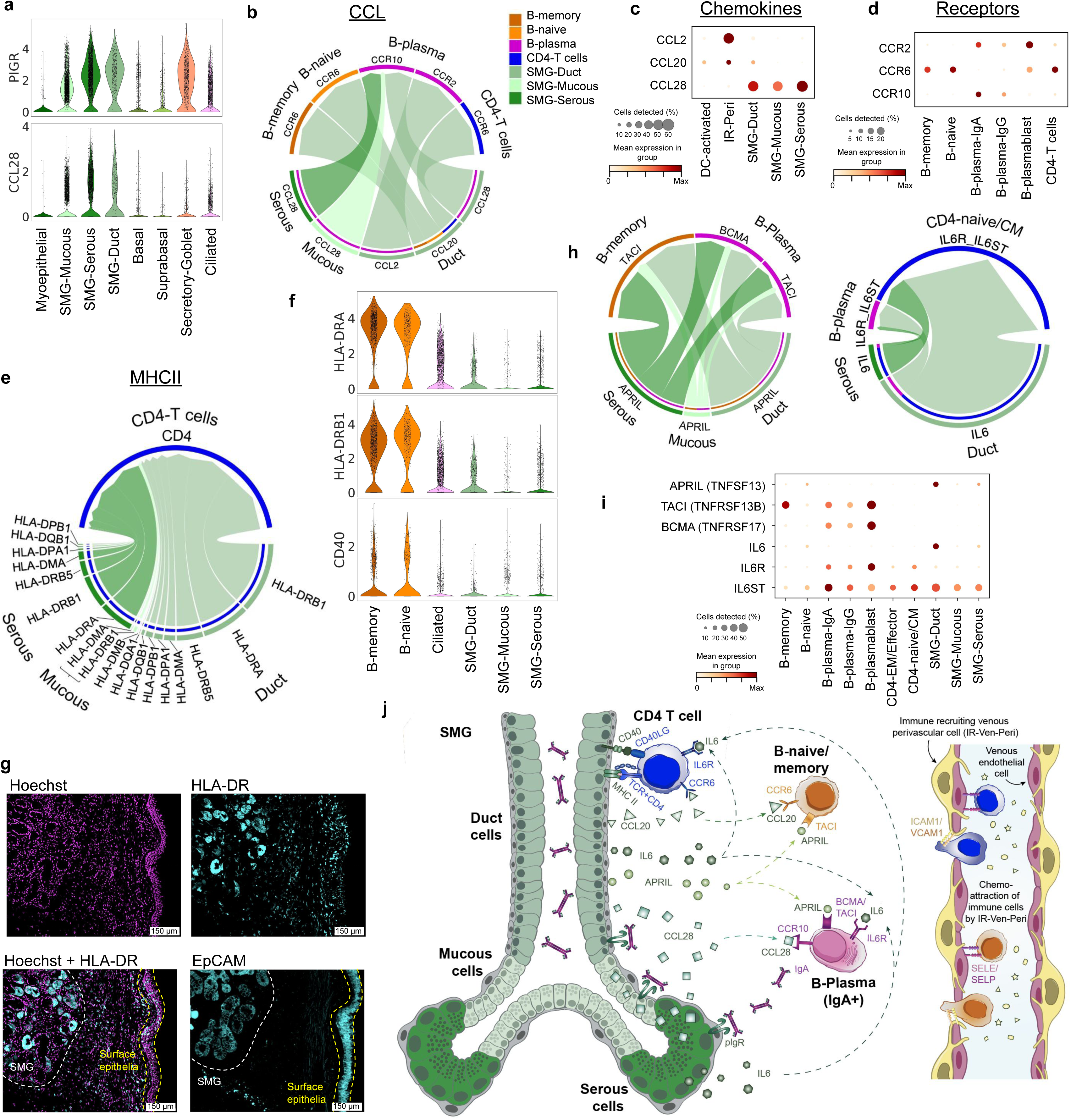
Cell-cell signaling at submucosal gland for B cell recruitment and survival niche. (a) Expression of PIGR and CCL28 in epithelial cells. (b) CellChat cell-cell interaction analysis pathways for CCL chemokines present from SMG epithelial cells to B cells (memory, naive and IgA/IgG/plasmablast combined) or CD4-T cells (CD4-naive/CM, CD4-EM/Effector and CD4-TRM combined) within airway tissue (trachea and bronchi). Arrow direction denotes chemokine-receptor pairs on specific cell types, arrowhead thickness reflects the relative expression of chemokine signal from each cell type. (c-d) Expression dot plot of relevant chemokines and corresponding receptors as shown in (b). (e) CellChat analysis as in (b) showing signalling from HLA genes expressed by SMG epithelial cells, signaling to CD4 on CD4-T cells. Arrow direction denotes ligand-receptor pairs on specific cell types, arrowhead thickness and proportion of the circle by each HLA gene/SMG epithelial cell type reflects the relative expression. (f) RNA expression of HLA-DRA, HLA-DRB1 and CD40 in B cells (as professional antigen presenting cell controls), ciliated and SMG epithelial cells from sc/snRNAseq. (g) Protein staining of HLA-DR and EpCAM in human trachea showing strong expression of HLA-DR in the SMG. (h-i) CellChat analysis as in (b) and (e) showing signalling of APRIL and IL-6 from SMG epithelial cells to relevant B cell subsets and CD4-naive/CM. Arrow direction denotes ligand-receptor pairs on specific cell types, arrowhead thickness and proportion of the circle for each gene/cell type reflects the relative expression. (i) Expression dot plot of relevant ligands and corresponding receptors as shown in (h). (j) Schematic of SMG immune niche showing immune cell recruitment and extravasation facilitated by venous endothelial cells and IR-Ven-Peri and signaling patterns between SMG gland epithelial cells, CD4-T cells, B-naive/memory cells and B-plasma cells to attract immune cells and promote antigen-specific T cell dependent and T cell independent pathways leading to IgA secretion at the SMG.

Cellchat also predicted interactions between *HLA* genes expressed by SMG epithelial cells, particularly SMG-Duct cells, and CD4-T cells (Figure 5e, f). The expression of *HLA-DRA* and *HLA-DRB1* in SMG-Duct cells was comparable to ciliated cells and, as expected, less than B memory and naive cells (Figure 5f). Interestingly, we observed strong expression of HLA-DR protein by IHC in the glands, but not in the surface epithelium showing discrepancy between transcript and protein levels (Figure 5g). We additionally found expression of *CD40,* a co-stimulatory molecule key for APC-T cell interactions, in SMG epithelial cells (Figure 5f). The expression of both these factors suggests that SMG-Duct cells can present antigen to CD4 T cells. Expression of HLA-DR and CD40 has previously been observed in airway and nasal epithelial cells, resulting in promotion of T cell proliferation *in vitro* (Rossi et al. 1990; Kalb et al. 1991; Cagnoni et al. 2004; Gormand et al. 1999; Tanaka et al. 2001). Interestingly, CD40 expression has been observed in duct, but not other cells, of the salivary gland and is upregulated in Sjögren’s syndrome, a systemic autoimmune exocrinopathy (Dimitriou et al. 2002).

In further support of an immune niche, A Proliferation Inducing Ligand (APRIL), a factor important for B cell survival, differentiation and class switching, was expressed by SMG-Duct cells, and to a lesser extent by SMG-Serous cells (Figure 5h,j). Cell-cell communication through these ligands was predicted between SMG epithelial cells and B-memory and B-plasma cells, based on differential expression of the receptors TACI and BCMA (Figure 5 h, j). In the colon, APRIL expression can be induced on intestinal epithelial cells and leads to IgA2 class-switch recombination (CSR) in the local tissue environment (He et al. 2007). We also find that a portion of B memory cells express activation-induced cytidine deaminase (*AIDCA*), suggesting the possibility of local CSR at the SMG through TACI-APRIL signaling (Figure S10 g). SMG-Duct, and to a lesser extent SMG-Serous, cells expressed *IL-6*, which was predicted to interact with *IL-6R/IL-6ST* on B plasma and a subset of CD4-T cells, the CD4-naive/CM T cells (Figure 5 i,j). In combination with APRIL, IL-6 induces and supports long lived plasma cells, and is a potent inducer of IgA secretion in IgA committed plasma cells (Beagley et al. 1989; Hirano et al. 1986). Furthermore, IL-6 has been shown as a required factor for CD4 T cell memory formation, and for overcoming Treg mediated suppression (Nish et al. 2014). Salivary gland epithelial cells have been shown to induce CD4 T cell differentiation into Tfh cells in an IL-6 dependent manner (Gong et al. 2014), which induced proliferation of B cells. In this study, proliferation of B cells was also induced in co-culture only with salivary gland epithelial cells, suggesting additional T-independent mechanisms. IL-6 is upregulated in serum and bronchoalveolar lavage fluid in asthma and COPD patients, suggesting that the balance of IL-6 signaling in the SMG is important for disease (Mercedes Rincon 2012; Savelikhina et al. 2018; Tillie-Leblond et al. 1999).

In conclusion, we have identified the localisation of IgA plasma cells, naive/memory B cells and T cells at the SMG along with a large number of molecular signalling pathways involved in B lymphocyte recruitment, antigen presentation and both T cell-dependent and -independent maturation and survival. These pathways are known to be functional in other secondary lymphoid organs, including within MALT, and we now show their possible involvement in establishing and maintaining the GALN (Figure 5k) of the lung.

## Summary and Conclusions

We have produced a detailed annotation of the transcriptomes of cell types in the human lung; assessing their different locations along the tracheobronchial tree and parenchyma. Within the over 193,000 high quality transcriptomes presented, we identify 77 cell types or states, including for the first time the transcriptomes of chondrocytes, peripheral nerve cells, and SMG duct cells as well as subtypes of Schwann cells, muscle and macrophages. Among the newly identified cell types we find specialised cell types expressing specific sets of chemokines within the fibroblast, smooth muscle and macrophage compartments. We also identify a novel IgA immune cell niche within the airway submucosal glands, which we term the gland associated lymphoid niche (GALN), with potential disease relevance.

The strength of this study comes from combining multiple complementary technologies: scRNAseq, scRNAseq, VDJ analysis and spatial transcriptomics. snRNAseq, though more limited in the number of genes detected than scRNAseq, is not biased by enzymatic treatments and dissociation artefacts. This has allowed us to define the transcriptomes of chondrocytes and perichondrial fibroblasts (PC-Fibro) and to separate rare cell types such as Schwann cell subsets and nerve-associated fibroblasts (NAFs). Visium ST has allowed us to demonstrate the localisation of cell types within the tissue; confirming those already known (such as ciliated epithelial cells), those previously transcriptionally undefined such as chondrocytes, and entirely undescribed cell types in the human lung.

We characterise cell types that may cooperate to generate an immune niche at the SMG, finding signalling circuits also observed in lymph nodes, MALT and immunologically active tissues such as the gut, resulting in both T cell-dependent and independent B cell responses. We propose cell-cell interactions between SMG epithelial cells, including the SMG-Duct cells, CD4-T cells and B-naive, -memory and IgA secreting plasma cells, that we map to the SMG. The survival, maturation and potentially class switching of B-lineage cells are supported by APRIL and IL-6, expressed by SMG-Duct cells, providing T cell independent factors. Additionally, the SMG epithelial cells have the potential capacity to induce T cell dependent immunity and B cell responses through expression of MHC-II and CD40. Further functional analysis will have to investigate our findings, examine the roles of the T cells we detect, and test the function of SMG-Duct cells as professional APCs and in promoting class switch recombination and somatic hypermutation at these sites.

Many of the signalling pathways we described have been observed in other tissues and are particularly well described for the recruitment of IgA plasma cells to the mammary gland and to the Peyer’s patches of the gut. No such secondary lymphoid structures or pathways within healthy lung tissue has been observed in the airways, but we postulate that the SMG microenvironment may fulfil a similar function.

Overall, the multi-omics data presented herein offer a comprehensive view of cell types and states within the human lung, with both macro- and micro-anatomical cell localisation information. We present new cell types with potential impact on a number of disease conditions both within and outside of the lung, and computationally analyse cellular interactions, advancing our understanding of lung composition and immunity. This comprehensive analysis has allowed us to take a systems approach that begins to define organ function as a result of the interactions between cells from distinct compartments. The interactions observed at the newly defined GALN may be relevant for other IgA secreting sites, such as mammary, salivary and tear glands.

Our findings are likely to be highly relevant to infectious disease biology, since IgA secretion is critical in combating infectious pathogens such as SARS-CoV-2 (Sterlin et al. 2021). We report higher proportions of IgA plasma cells in the airways of COVID-19 patients, supporting the importance of IgA immune niche. Nasal vaccines could induce a strong local sIgA response, yet all currently licenced COVID-19 vaccines are administered intramuscularly (Lund and Randall 2021). These vaccines are highly effective in preventing serious disease, but preclinical studies suggest that they provide little protection against viral replication and shedding in the upper airway (Bleier, Ramanathan, and Lane 2021; van Doremalen et al. 2021). Novel nasal vaccines are being developed that in non-human primates have shown promising prevention of replication and shedding of the virus due to the induction of a mucosal IgA response in the upper and lower respiratory tract (reviewed in (Tiboni, Casettari, and Illum 2021)). Whether the same is true in humans remains to be determined, but in COVID-19 patients, SARS-CoV-2 neutralisation was more closely correlated with IgA than IgM or IgG (Sterlin et al. 2021), and increased titres of IgA were observed in tears, nasal fluid and saliva (Cervia et al., n.d.) further highlighting the importance of the IgA immune niche in COVID-19 immunity. A better understanding of the mucosal IgA immune niche and the pathways that establish this niche, which we describe here, may offer options to augment this immune response.

## Acknowledgements

We thank Jana Eliasova for the graphical illustrations. Chuan Xu provided a script for TCR gene usage analysis. Cecilia Dominguez Conde supported with a script for TCR clonotype sharing analysis. Martin Prete’s and the Cellgen IT computational support has been central for the analysis. We are grateful to the organ donors, their families and the Cambridge Biorepository for Translational Medicine for the gift of human tissue.

K.B.M. and S.A.T. are supported by Wellcome (WT211276/Z/18/Z and Sanger core grant WT206194). E.M. is supported by ESPOD fellowship of EMBL-EBI and Sanger Institute. The project has received funding from the European Union’s Horizon 2020 research and innovation programme under grant agreement No 874656. This project has been made possible in part by grant 2019-202654 from the Chan Zuckerberg Foundation.

## Author Contributions

K.B.M and E.M. conceived and designed the experiments; E.M., A.J.O., K.P., A.R.O., J.P.P., and N.H. carried out computational analysis; V.K. ran and optimised Cell2Location analysis.; A.W-C. helped with experimental planning, sample management and spatial gene expression; L.M., L.B., A.K., E.P., A.H., and A.O. carried out tissue dissociation and sc and snRNAseq experiments; M.D., L.T., S.P. and S.F.V. performed Visium Spatial Transcriptomics and RNAScope analysis, supervised by M.P; N.R. carried out IHC and protein staining; P.H. and R.E. contributed to cell types annotation; K.M., N.G., K.S-P provided human tissue samples; N.K. carried out statistical analysis; O.A.B., M.C., O.S., S.A.T. and K.B.M provided funding, discussion and supervision; and E.M., A.J.O., A.W-C and K.B.M wrote the manuscript.

## Data availability

Sequencing data for scRNAseq, snRNAseq, VDJ-seq and Visium ST will be available via the DCP (http://data.humancellatlas.org). Sc/snRNAseq data will be available for browsing and gene expression data download via cellxgene website.

## Supplementary legends

Figure S1. Overview of human lung dataset across five locations (a) H&E sections of the deep human tissue biopsies from multiple regions showing all major structures of the lungs and airways. (b) Expression of cell type marker genes in the master cell type groups, from both single cell and single nuclei RNAseq combined. (c) Protein staining of chondrocyte markers in the cartilage of human bronchus from the Human Protein Atlas. (d) Proportion of mesenchyme cell type groups in the airways from cells and nuclei. (e) UMAP of sequencing material (cells or nuclei) and location (trachea, bronchi, parenchyma). (f) Variance of gene expression explained by metadata variables in the combined sc/snRNA-seq dataset, scRNA-seq and snRNA-seq datasets.

Figure S2. Overview of Spatial Transcriptomics slides used in the study. H&E staining as well as the number of UMI counts per spot are visualised for each section. RNA-seq sample names match those in Supplementary Table 3.

Figure S3: Novel fibroblast subsets (a) Dot plot of marker gene expression for indicated cell types. (b) UMAP of location and sequencing material from fibroblasts. (c) Heatmap showing annotated cell types to the predicted labels for fibroblasts from Travaglini et al. (Travaglini et al. 2020) by the Azimuth tool, colored by proportion. Labels by the proportion of annotated cells and the total number of cells mapping to the reference. (d) Violin plots with predicted annotation score for each of the annotated cell types to the reference. Small dots represent cells, circles represent mean values and bars show standard deviation.

Figure S4: Validation of immune recruiting fibroblasts and their tissue localisation. (a) smFISH staining in human bronchi tissue for IR-Fibro markers (CCL21, CCL19) showing independent localisation from immune cells (PTPR) and smooth muscle cells (ACTA2). (b) H&E staining on Visium ST with manually annotated regions for the immune infiltrate in blue. Cell2location mapping density scores with zoom into the region of interest, showing density values for IR-Fibro and relevant immune cells from the current lung study as well as for germinal center cell types from a gut dataset (R. Elmentaite et al. 2021). Dashed lines are added for better visual comparison between the cell types and regions.

Figure S5: Peribronchial and perichondrial fibroblasts. (a) Protein staining of PB-Fibro markers (COL15A and ENTPD1) in human bronchus sections from the Human Protein Atlas. (b) Marker genes for PB-fibro and PC-fibro. (c) Protein staining of PC-Fibro marker (COL12A1) in human bronchus from the Human Protein Atlas mapping to cartilage. (d) UMAP of adventitial fibroblasts, PC-fibro and chondrocytes from single nuclei data colored by monocle 3 pseudotime and cell type. (e) Expression of genes associated with bone/cartilage function, markers of PC-fibro and cartilage genes in the nuclei as shown on (d), ordered by pseudotime. (f) PC-fibro marker gene enrichment in Human Phenotype Ontology by g:Profiler.

Figure S6. Schwann cells and nerve-associated fibroblasts (NAF). (a) Marker dot plot for myelinating, non-myelinating Scwhann cells and for epi- and endoneurial NAF-s. (b) g:Profiler enrichment results for myelinating Schwann cell markers with detailed results for myelination and transcription factor EVX1. (c) g:Profiler enrichment results for non-myelinating Schwann cell markers. (d) Expression in Transcript per million (TPM) of NAF markers in GTEx bulk RNA-seq data. (e) Visium ST H&E staining of human bronchi, with enlarged nerve bundle and Cell2location cell type mapping density scores for Schwann and NAF cell types. (f-h) Human Protein Atlas antibody staining of (f) non-myelinating Schwann cell markers (CADM, GRIK2, NCAM1, ITGB4 and L1CAM) (g) endoneurial NAF marker (USP54) and (h) epineurial NAF markers (SLC22A3 and SORBS1) within the nerve bundles in human bronchus. Arrows indicate nerve bundles. (i) RNAscope staining for myelinating (MLIP) and non-myelinating (SCN7A, SOX10) Schwann cell and epineurial (SLC2A1) NAF specific genes in bronchial nerves. (j) Expression of neuropathy associated genes in Schwann and NAF cell types. Previously unknown cell type specific expression shown in color: light green for novel expression pattern, light blue for distinguishing expression for nmSchwann cells.

Figure S7. Vascular and smooth muscle cell types. (a) Markers dot plot for vascular endothelia. (b) Bronchi section with H&E and Cell2location analysis density score for airway smooth muscle population on a Visium ST slide. (c) NPR2 staining in oesophagus and bronchus from the Human Protein Atlas. Black arrows indicate the airway and oesophagus surrounding non-vascular smooth muscle. (d) ASM marker expression in all GTEX tissues. Tissues are ordered by unsupervised clustering based on expression similarity. The dotted line highlights tissues which are surrounded by a thick smooth muscle layer. The orange rectangle shows muscular tissues from oesophagus, and the blue rectangle shows the non-muscular mucous layer of oesophagus tissue. (e) Cell2location density scores of pulmonary and vascular endothelium for parenchyma and bronchi Visium ST sections. (f) IR-Ven-peri markers localise at the venous vessels in the airway. smFISH staining for IR-Ven-peri (CCL21, CCL19), venous endothelia (ACKR1) and smooth muscle (ACTA2) markers. (g) Leukocyte rolling and homing genes, and chemokines expressed in Endothelia and Perivascular cells together with their interaction partners expression in immune cell groups. Interaction partners are indicated with blue shades.

Figure S8. Epithelial cell annotations and location specific ciliated cell gene expression. (a) Marker gene expression dot plot for epithelial cells. (b) UMAP of epithelial cells (excluding AT1 and AT2 cells) from scRNAseq data. (c) RNA velocity results on UMAP plot for single cells from airway epithelia, with colors indicating cell types as in (b). (d) Cell type proportion analysis with fold changes and Local True Sign Rate (LTRS) score for all cell type groups with regards to location, donor, sequencing material and dissociation protocol. (e) smFISH staining for mucous (MUC5B), serous (LPO) and duct (MIA / ALDH1A3 / RARRES1) cell markers in human bronchus section. (f) smFISH staining of secretory goblet/club (SCGB1A1), ciliated (FOXJ1) and duct (ALDH1A3 / RARRES1) cells in human bronchus section. (g) smFISH staining for muscle (TAGLN), basal epithelia (KRT14), duct (ALDH1A3) and myoepithelium (FHOD3) cell type markers in human bronchus section. (h) H&E from bronchial section and Cell2location density values for mapping duct, mucous, serous, ciliated and myoepithelial cells onto the Visium ST section.(i) Linear mixed model analysis revealed 80 differentially expressed genes in ciliated cells between trachea and other locations (bronchi and parenchyma). Violin plots of normalised log-transformed expression separated by location in the single nuclei RNA-seq data for three representative genes upregulated in nasopharyngeal carcinoma gene set from GSEA database with LTSR>0.9 consistently higher expressed in the trachea. (j) SARS-CoV-2 receptor and viral entry gene expression in airway epithelial cells, with expression levels of ACE-2 in the ciliated cells from snRNAseq data shown by location in a violin plot.

Figure S9. Immune cell type groups. (a) Marker genes dot plot for myeloid cells. (b) Marker genes dot plot for T&NK cells. (c) UMAP for Megakaryocytes and Mast cells along with marker gene expression dot plot. (d) Cell2Location density scores for CD8-TRM, Ciliated and CD8-EM cell types in human bronchi sections and corresponding H&E. (e) Fraction of clonally expanded cells in T & NKT cell types from VDJ data. (f) Proportion of shared TCR clonotypes between samples from VDJ data. Color bars indicate location and donor. (g) Effect of location, donor, material and protocol on immune cell type proportions. Cell type proportion analysis with fold changes and Local True Sign Rate (LTRS) score for myeloid cell types, T & NK cell types and B lineage cell types.

Figure S10. Additional B cell data (a) Marker gene expression dot plot for B-lineage cells. (c, d) Human Protein Atlas staining for B plasma marker MZB1 in the bronchus (c) nasopharyngeal glands (d). (e) Number of B lineage cells with different Ig isotypes in parenchyma from the analysis of VDJ amplified libraries. (f) Multiplex IHC of human trachea for Ig isotypes showing distinction between glands and non-gland regions of tissue (g) Violin plot for expression of AICDA in B cell subsets.

Supplementary File 1. Cell type marker genes.

Marker genes for all the described cell types.

Supplementary Table 1. Sample and donor info for single-cell RNA-seq.

Sample ID-s and corresponding location, spatial code, material of cells/nuclei, protocol, enrichment and dissociation notes, Donor ID-s with age, BMI, gender and smoking history if available, 10x version, Gene expression sequencing run ID and corresponding BCR and TCR sample ID-s for single cell RNA-seq samples.

Supplementary Table 2. Sample info for single-nuclei RNA-seq.

Sample ID-s and corresponding information about pooling, donors, location, protocol and 10x version for single nuclei RNA-seq samples.

Supplementary Table 3. Sample info for Visium Spatial Transcriptomics.

Sample and image ID-s, location and donor information, permeabilization time and Visium slide ID for Visium ST samples.

Supplementary Table 4. Manual annotation of tissue regions on Visium ST. Manual annotation indicates presence (y) of various tissue regions on every Visium ST samples: Airway Smooth Muscle, Arterial vessel, Cartilage, Glands, Mesothelium, Multilayer epithelium, Nerve, Parenchyma, Perichondrium, Pulmonary vessel, Small airway, Venous vessel, iBALT-like immune infiltrate. The annotations on the tissue regions can be seen as categories on the accompanying loupe files stored in DCP.

Supplementary Table 5. Antibody information for IBEX staining. Staining cycle, antibody protein names, clone, conjugate, Vendor, catalog number and dilution used.

## Methods

### Aim and study design

We aimed to generate a multi-omics view of different regions of the human lung using single cell and nuclei sequencing (10X Genomics droplet sequencing) and spatial analysis (Visium Spatial Transcriptomics and RNAScope in situ hybridisation).

### Experimental Methods

#### Access to human tissue

Samples were obtained from deceased transplant organ donors by the Cambridge Biorepository for Translational Medicine (CBTM) with informed consent from the donor families and approval from the NRES Committee of East of England – Cambridge South (15/EE/0152). This consent includes generation of open-access genetic sequencing data and publication in open access journals in line with Wellcome Trust policy. CBTM operates in accordance with UK Human Tissue Authority guidelines.

#### Tissue dissociation and single cell sequencing

Tissue was collected from seven donors from five lung locations including trachea, bronchi at the 2nd/3rd generation, bronchi at the 4th generation, upper left lobe parenchyma and lower left lobe parenchyma (Figure 1a and Supplementary Tables 1-3). Following collection at the clinic, samples (ranging 1-4cm^3^) were immediately placed into cold Hypothermasol FRS as per the protocol outlined by Madissoon et al. (Madissoon et al. 2019) and transferred to the processing laboratory. Within 12h of cessation of circulation samples were divided into fresh portions of around 1g for immediate dissociation and pieces for embedding in OCT / freezing in isopentane cooled on dry ice for later spatial analysis and nuclei isolation. The majority of samples (n=5) were digested using liberase and trypsin, and CD45 positive cells loaded on 10X as a separate fraction (protocols.io 39ygr7w), although one donor was digested with collagenase D for comparison (protocols.io 34kgquw). Details on which samples received which treatments can be found in Supplementary Table 1&2; all protocols are available on protocols.io and linked in the metadata collated in the Human Cell Atlas Data Co-ordination Platform. Briefly, for the main tissue dissociation protocol used (for 5 donors), a 1g piece of lung tissue was washed with PBS-/- and minced finely with scalpels, before treatment with 13U/ml liberase TL and 0.1mg/ml DNase I for 30 minutes at 37°C with rocking. Cells were filtered through a 70μm strainer, washed with neutralisation media (RPMI + 20% FBS) and pelleted (sample P1). Tissue remaining in the cell strainer was further digested with 0.25% trypsin-EDTA with DNase I for 30 minutes at 37°C with rocking, filtered and washed with neutralisation media. Meanwhile, sample P1 was treated with red blood cell lysis buffer before being separated into CD45 positive and negative fractions using MACS (Miltenyi, as according to the manufacturer’s protocol). The CD45 negative fraction was pooled with cells from trypsin treatment, resulting in two samples for loading on 10X: CD45 positive cells from liberase TL digestion (to enrich for immune cells) and pooled CD45 negative liberase-treated cells with trypsin-treated cells (non-immune fraction). These two cell fractions were resuspended in 0.04% BSA / PBS, counted using a Nucleocounter (Chemometec) and loaded on the 10X Genomics Chromium instrument for single cell sequencing, aiming to capture 5000 cells. The majority of samples were sequenced using 10X Genomics 5’v1 chemistry (5 donors; 2 were donors run using 3’v2 chemistry; Supplementary Table 1) according to the manufacturer’s protocol.

#### Single nucleus sequencing

Single nuclei were isolated from frozen tissue using a modified version of the protocol described by Krishnaswami et al., 2016. Frozen tissue sections (8 50 um thick sections) were homogenised using a glass dounce homogeniser (Sigma) in Nuclei Isolation buffer (NIM - Sucrose 0.25 M, MgCl2 0.005 M, KCl 0.025 M, Tris (buffer pH7.4) 0.01 M, DTT 0.001 M and Triton X-100 0.1%) in the presence of Complete protease inhibitors (Roche) and RNAse inhibitors RNasin (Promega) – 0.4 U/ul and Superasin (Invitrogen) 0.2 U/ul). Tissue was homogenised using ∼15 strokes with pestle A (clearance 0.0028-0.0047 in.) then pestle B (clearance 0.0008-0.0022 in.). Isolated nuclei were filtered through a 40 uM filter, collected at 2000 x g followed by resuspension in 0.5 ml of storage buffer (PBS containing 4% BSA and RNasin (Promega) – 0.2 U/ul. Nuclei were incubated with NucBlue® (ThermoFisher), 1 drop was added to 0.5 ml nuclei suspension. Nuclei were purified by FACs sorting, stained with Trypan blue and counted using a disposable haemocytometer. Nuclei were diluted to an appropriate concentration and 5,000 nuclei from 5 different samples were pooled and analysed using the 10X 3’ platform. 25,000 nuclei were and loaded onto the 10X Chromium controller using the 3’ v3.1 kit as per the Chromium Single Cell 3ʹ Reagent Kits v3 User Guide, targeting to recover ∼3,000 nuclei per sample. Post GEM-RT cleanup, cDNA amplification and 3’ gene expression library construction were carried out according to the appropriate user guide. The resulting libraries were sequenced on the Novaseq platform.

#### Visium Spatial Transcriptomics

Samples no larger than 0.5cm^2^ were cut from each of the 5 locations of lung (trachea, bronchi at the 2nd/3rd generation, bronchi at the 4th generation, upper left lobe parenchyma and lower left lobe parenchyma) and additional small airway. Majority of the parenchyma was removed from the airway tissues, embedded in OCT and flash frozen in dry-ice cooled isopentane at -55°C. Haematoxylin and eosin staining was used to determine the morphology of tissue blocks, in order to choose those to proceed to Visium. Ten micron sections were then cut from the blocks onto Visium slides (10X Genomics) and processed according to the manufacturer’s protocol. Full list of samples, locations, donors and permeabilisation times are shown in Supplementary Table 3 and Supplementary Figure 3. Haematoxylin and eosin images generated during the Visium protocol were captured at 20x magnification on a Hamamatsu Nanozoomer S60 and exported as tiled tiffs for analysis.

Dual-indexed libraries were prepared as in the 10X Genomics protocol, pooled at 2.25nM and sequenced 4 samples per Illumina Novaseq SP flow cell with read lengths 28bp R1, 10bp i7 index, 10bp i5 index, 90bp R2.

#### RNAScope

Tissue blocks for RNAScope in situ hybridisation were chosen based on haematoxylin and eosin staining results as described under ‘Visium’. Ten micron-thick cryosections cut on to superfrost plus slides were processed using the RNAScope 2.5 LS multiplex fluorescent assay (ACD, Bio-Techne) on the Leica BOND RX system (Leica). Fresh frozen lung sections were fixed for 90 minutes with chilled 4% paraformaldehyde, washed twice with PBS and dehydrated through an ethanol series (50%, 70%, 100%, 100% ethanol) before processing according to the manufacturer’s protocol with protease IV treatment. Samples were first tested with RNAScope positive and negative control probes before proceeding to run probes of interest. Slides were stained for dapi (nuclei) and three (or four where indicated) probes of interest, with fluorophores opal 520, opal 570, opal 650 and atto 425 (4-plex only) at between 1:500 - 1:1000 concentration. These were then imaged on a Perkin Elmer Opera Phenix High Content Screening System with water immersion at 20x Magnification.

#### Tissue preservation and antibody staining

The fresh airway tissue was received in cold Hypothermasol, fixed with 1% PFA solution for 24h at 4°C and transferred to cold 30% sucrose solution overnight, before freezing in OCT. The fixed tissue was sectioned at 10µm thickness. Iterative staining of human trachea sections was performed as described by Radtke et al. (Radtke et al. 2020). 10µm sections were permeabilised and blocked in 0.1M TRIS, containing 0.1% Triton (Sigma), 1% normal mouse serum, 1% normal goat serum and 1% BSA (R and D). Samples were stained for 2h at RT (primary antibodies) or 1h at RT (secondary antibodies) in a wet chamber with the appropriate antibodies, washed 3 times in PBS and mounted in Fluoromount-G® (Southern Biotech). Images were acquired using a TCS SP8 (Leica, Milton Keynes, UK) inverted confocal microscope. The coverslip was then removed and slides were washed 3 times in PBS to remove any mounting medium. Bleaching of the fluorochromes was achieved using a 1mg/mL solution of lithium borohydride in water (Acros Organics) for 15 minutes at RT. The slides were then washed 3 times in PBS prior to staining with a different set of antibodies as described above. The process was repeated a total of 5 times. Raw imaging data were processed using Imaris (Bitplane) using Hoechst as fiducial for the alignment of subsequent images. The staining set-up and antibody information is in Supplementary Table 5.

### Computational analysis

#### Mapping of Gene expression libraries

Single-cell and single nuclei RNA-seq gene expression libraries were mapped with Cell Ranger 3.0.2, and Visium libraries were mapped with Space Ranger 1.1.0 from 10x genomics (https://support.10xgenomics.com). Both types of libraries were mapped to an Ensembl 93 based reference (10x-provided GRCh38 reference, version 3.0.0). For nuclei samples, the reference was altered into a pre-mRNA reference as per 10x instructions. TCR/BCR libraries were mapped with Cell Ranger 4.0.0 to the 10x-provided VDJ reference, version 4.0.0.

#### Spatial mapping of cell types using Visium Spatial Transcriptomics and cell2location

Visium Spatial Transcriptomics (ST) data was used to establish tissue locations of the cell types presented in the paper. This was achieved by applying the Cell2location method to estimate the abundance of each cell type in the Visium slides by integrating the single cell/single nuclei and spatial transcriptomes (Kleshchevnikov et al., n.d.). The first step of cell2location workflow is estimation of reference expression signatures of cell types representing gene expression profiles for each annotated cell type and subpopulation (n=85). This step was performed using Negative Binomial regression (from cell2location package) that allows accounting for technology (sc/sn), batch and donor effects. To spatially map iBALT-like immune infiltrate regions we extended the cell type reference by including immune populations involved in the germinal center reaction. This was done by extending the above mentioned set of reference cell type signatures with the signature of these populations which were derived using published human gut data sets that featured good representation of such tissue infiltrating populations (R. Elmentaite et al. 2021). In the second step of cell2location workflow these reference signatures are used to estimate absolute spatial abundance of cell types, integrating and normalising data across 11 Visium sections (5 were excluded based on quality control metrics). 10X Visium data was processed as described above to untransformed and unnormalised mRNA counts, filtered to genes shared with scRNA-seq, which were used as input to cell2location with hyperparameters that were determined using information about the tissue and experiment quality:

1. Expected cell abundance per location 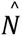 = 20, which were estimated by computing average cell count across 20 representative locations derived by manually counting cells in histology images.
2. Regularization of within-experiment variation in RNA detection sensitivity α^y^ = 20, representing more lenient regularisation to account for large technical within-slide variability in total count observed in this data.

The mode was trained until convergence was achieved (40,000 iterations).

To aid interpretation, all locations in the Visium data were annotated with anatomical regions label using paired histology images. Each location on each Visium slide that was aligned with the tissue was annotated as “Tissue” (manual annotation) and was used for analysis. Specific tissue regions were annotated as follows: Cartilage, Glands, Multilayer epithelium, Parenchyma, Nerve bundle, Perichondrium, Airway smooth muscle, Arterial vessel, Venous vessel, Weird morphology, Mesothelium, Pulmonary vessel, Immune infiltrate (iBALT-like), Small airway (on parenchyma sections only). The annotation was used to compute average estimated cell abundance of each cell type across annotated regions, which was normalised per cell type and presented as a dotplot.

#### Single cell and nuclei RNA-seq analysis

The CellRanger unfiltered matrices were used as an input for the SoupX algorithm (M. D. Young and Behjati 2020) to remove ambient RNA contamination. The single cell RNA-seq libraries were corrected according to the tutorial (https://github.com/constantAmateur/SoupX). For each single nuclei RNA-seq library, CellRanger filtered nuclei were first subjected to a set of QC filters. Pass QC nuclei were processed using standard scanpy pipeline and were clustered at a relatively low resolution to form 5-10 clusters. Nuclei that didn’t pass QC were assigned to those clusters by logistic regression. This clustering was then passed to SoupX, which used it to derive a set of cluster-specific genes for automatic estimation of contamination rate, which together with estimated soup composition were used by SoupX to clean the data. Default values were used when calling SoupX’s functions, except for “autoEstCont()” where “soupQuantile” was set to 0.8 to improve stability of estimation. The single-cell and nuclei libraries with SoupX correction were analysed using the standard scanpy 1.7.1 workflow (F. A. Wolf, Angerer, and Theis 2018). The cells with >4000 counts in nuclei and 20,000 counts in the cells were removed. In the cells, droplets with >10,000 features were removed. Lower threshold of 1000 features was applied to donor A37 due to ambient RNA contamination that was difficult to remove. The rough level clustering was used to annotate master cell types. Master cell types were further subtracted and re-analysed with scanpy workflow including new highly variable genes detection. Typically, between 1000 and 3000 highly variable genes (HVG-s) were used for calculating 40 principal components that were used for calculating the UMAP coordinates. Data integration via Harmony (Korsunsky et al. 2019) or BBKNN (Polański et al. 2020) or scVI (Lopez et al. 2018) was used with either “Material” as input to correct between cells and nuclei, or “Material and Donor” in order to take into account the variability between donors. Clustering at different resolutions was done in order to determine biologically meaningful cell types. Doublets and contaminated cells were detected via overclustering and observing markers from more than one cell type, and in case of nuclei-seq also observing higher counts. Doublet clusters were removed. Iterative clustering approach was used to further derive clusters for less abundant cell types. In addition to known marker genes, new ones were derived using the scanpy’ rank_genes_groups function. Cell types and master clusters were annotated according to known and newly derived markers as in Supplementary File 1.

#### Post-processing analysis

Gene set enrichment analysis was performed in GSEA online tool (https://www.gsea-msigdb.org/gsea/index.jsp)(Subramanian et al. 2005) for specific gene sets and in gProfileR (https://biit.cs.ut.ee/gprofiler/gost). The analysis of gene expression in GTEx tissues was performed in the GTEx portal (https://gtexportal.org/home/). Cell-cell interaction analysis was performed with CellPhoneDB (https://www.cellphonedb.org/) (Efremova et al. 2020) and with CellChat (http://www.cellchat.org/). In order to cope with donor to donor differences, the dataset was downsampled to a set number of cells per every donor-celltype combination.

Pseudotime analysis for selected cell populations was performed with Monocle3 (ref), its functionality to infer a pseudotime based on UMAP coordinates. The root was identified as the cell with the highest combined expression of canonical progenitor markers (VCAN for chondrocytes; TGM2, HMCN2 and SULF1 for smooth muscle). Cell trajectory analysis was performed using the scVelo package (v0.2.1)(Bergen et al. 2020) and specifying stochastic model.

Label transfer was performed via Azimuth tool (https://azimuth.hubmapconsortium.org/) with the lung reference data (Travaglini et al. 2020). Human protein atlas (https://www.proteinatlas.org/humanproteome/tissue) was used for extracting images of protein stainings with antibodies on human tissues.

#### BCR and TCR analysis from VDJ-data

VDJ analysis was done with Scirpy 0.6.0 (https://icbi-lab.github.io/scirpy/)(Sturm et al. 2020). For TCR data, clonotypes were defined based on CDR3 nucleotide sequence identity. For BCR data, clonotypes were defined based on the Hamming distance between CDR3 amino acid sequences with a cutoff of 2 and orphan VJ chains removed. In both cases, V gene identity was required and the CDR3 sequence similarity was evaluated across all of a cell pair’s V(D)J chains.

#### Poisson linear mixed model for cell type composition analysis

Poisson regression with various metadata as covariates was applied to adjust confounding effects on the cell type count data as previously described (Stephenson et al. 2021; Rasa Elmentaite et al. 2020). We used Location as biological factor and Protocol, Material (scRNAseq vs snRNAseq) and Donor as technical factors in the model as random effects to overcome the collinearity (see Supplementary Notes in Elmentaite et al. for more details).

#### Variance in gene expression

In order to determine the effects of the metadata features on the expression data, a linear mixed model was used (A. M. H. Young et al. 2021). Genes expressed in less than 5% of the samples were filtered out. The count matrix was then normalised and log-transformed. The percentages of variance in gene expression data explained by each metadata feature were obtained through fitting the linear mixed model. The Bayes factor was then computed to determine the gene-specific effects of some metadata features in the expression data, assigning an effect size and a local true sign rate for all genes analysed. Genes presenting a local true sign rate (ltsr) value greater than 0.9 were considered significantly affected by the metadata feature analysed. See Supplementary Notes of (A. M. H. Young et al. 2021) for more details.

#### Functional GWAS analysis (fGWAS)

The fGWAS approach to determine disease relevant cell types is described elsewhere (R. Elmentaite et al. 2021). Summary statistics for the selected GWAS study of COPD (Shrine et al., 2019) were obtained via Open Targets Genetics (https://genetics.opentargets.io/study/GCST007431; https://www.ebi.ac.uk/gwas/studies/GCST007431).

#### Cell-cell interaction analysis

Cell-cell interaction from scRNAseq data was predicted using CellChat (Jin et al. 2021). B plasma subsets were combined (B-plasma) and cell types of interest (B-naive, B-memory, SMG-Duct, SMG-Mucous, SMG-Serous, CD4-naive/CM, CD4-EM/Effector and CD4-TRM) were downsampled to 200 cells per cell type per donor. Analyses were performed both with individual CD4-T cell subsets and all CD4 subsets combined. Normalised count matrix along with cell annotation metadata was processed through the standard CellChat pipeline, except that the communication probability was calculated with a truncated mean of 10%.

