## Supplementary Figures for "A spatial multi-omics atlas of the human lung reveals a novel immune cell survival niche"

Figure S1

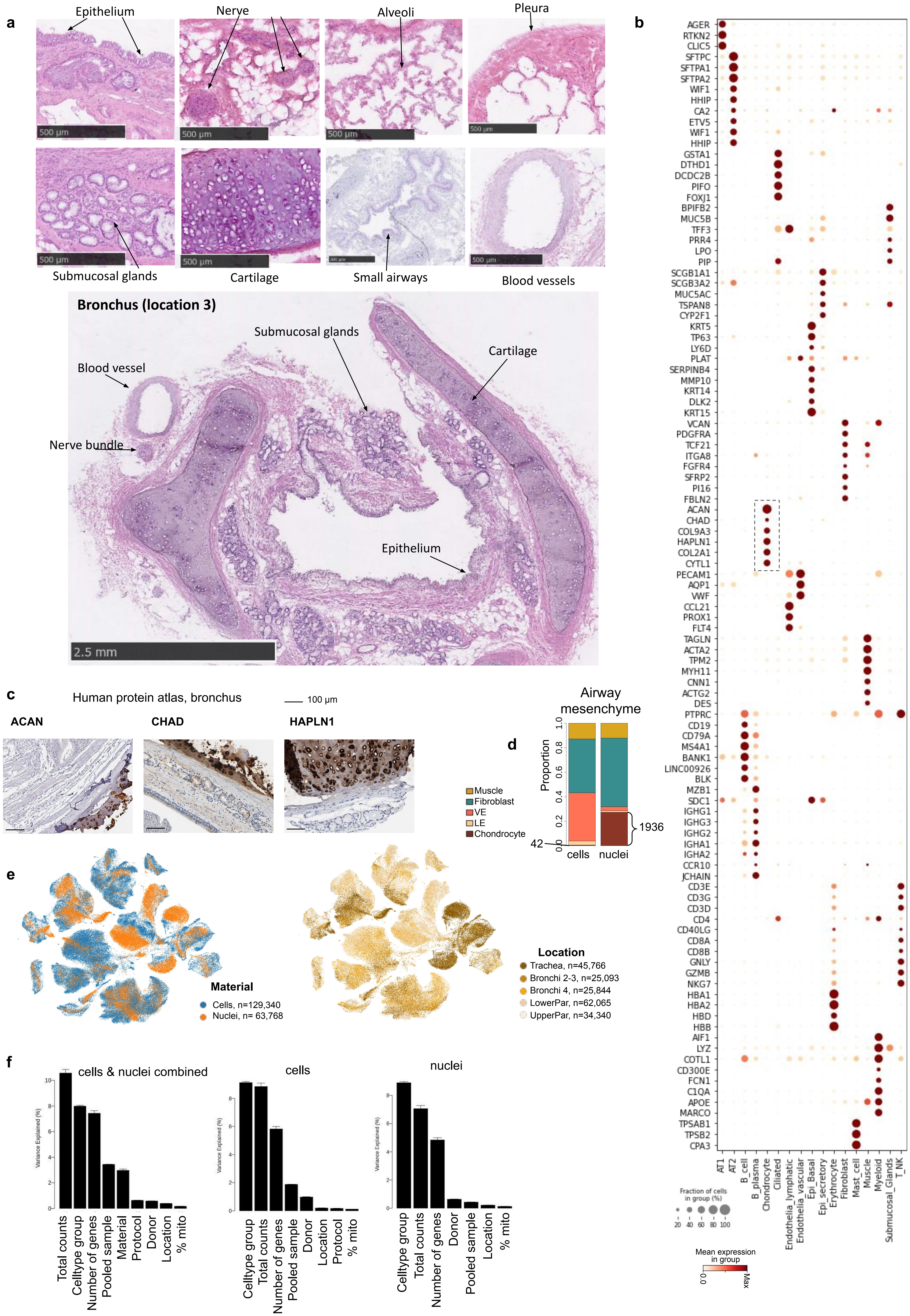
a
Epithelium
Nerve
Alveoli
Pleura
500 µm
Submucosal glands
Cartilage
Small airways
Blood vessels
Bronchus (location 3)
Submucosal glands
Blood vessel
Cartilage
Nerve bundle
Epithelium
2.5 mm
b
AGER
RTKN2
CLIC5
SFTPC
SFTPA1
SFTPA2
WIF1
HHIP
CA2
ETV5
WIF1
HHIP
GSTA1
DTHD1
DCDC2B
PIFO
FOXJ1
BPIFB2
MUC5B
TFF3
PRR4
LPO
PIP
SCGB1A1
SCGB3A2
MUC5AC
TSPAN8
CYP2F1
KRT5
TP63
LY6D
PLAT
SERPINB4
MMP10
KRT14
DLK2
KRT15
VCAN
PDGFRA
TCF21
ITGA8
FGFR4
SFRP2
PI16
FBLN2
ACAN
CHAD
COL9A3
HAPLN1
COL2A1
CYTL1
PECAM1
AQP1
VWF
CCL21
PROX1
FLT4
TAGLN
ACTA2
TPM2
MYH11
CNN1
ACTG2
DES
PTPRC
CD19
CD79A
MS4A1
BANK1
LINC00926
BLK
MZB1
SDC1
IGHG1
IGHG3
IGHG2
IGHA1
IGHA2
CCR10
JCHAIN
CD3E
CD3G
CD3D
CD4
CD40LG
CD8A
CD8B
GNLY
GZMB
NKG7
HBA1
HBA2
HBD
HBB
AIF1
LYZ
COTL1
CD300E
FCN1
C1QA
APOE
MARCO
TPSAB1
TPSB2
CPA3
AT1
AT2
B_cell
B_plasma
Chondrocyte
Ciliated
Endothelia_lymphatic
Endothelia_vascular
Epi_Basal
Epi_secretory
Erythrocyte
Fibroblast
Mast_cell
Muscle
Myeloid
Submucosal_Glands
T_NK
Fraction of cells in group (%)
20 40 60 80 100
Mean expression in group
0.0
Max
c
Human protein atlas, bronchus
100 µm
ACAN
CHAD
HAPLN1
d
Airway mesenchyme
Proportion
0.0
0.2
0.4
0.6
0.8
1.0
Muscle
Fibroblast
VE
LE
Chondrocyte
42
1936
cells
nuclei
e
Material
Cells, n=129,340
Nuclei, n= 63,768
Location
Trachea, n=45,766
Bronchi 2-3, n=25,093
Bronchi 4, n=25,844
LowerPar, n=62,065
UpperPar, n=34,340
f
cells & nuclei combined
Variance Explained (%)
0
2
4
6
8
10
Total counts
Celltype group
Number of genes
Pooled sample
Material
Protocol
Donor
Location
% mito
cells
Celltype group
Total counts
Number of genes
Pooled sample
Donor
Location
Protocol
% mito
nuclei
Celltype group
Total counts
Number of genes
Donor
Pooled sample
Location
% mito

Figure S2

### H&E, number of total UMI for all Visium ST sections

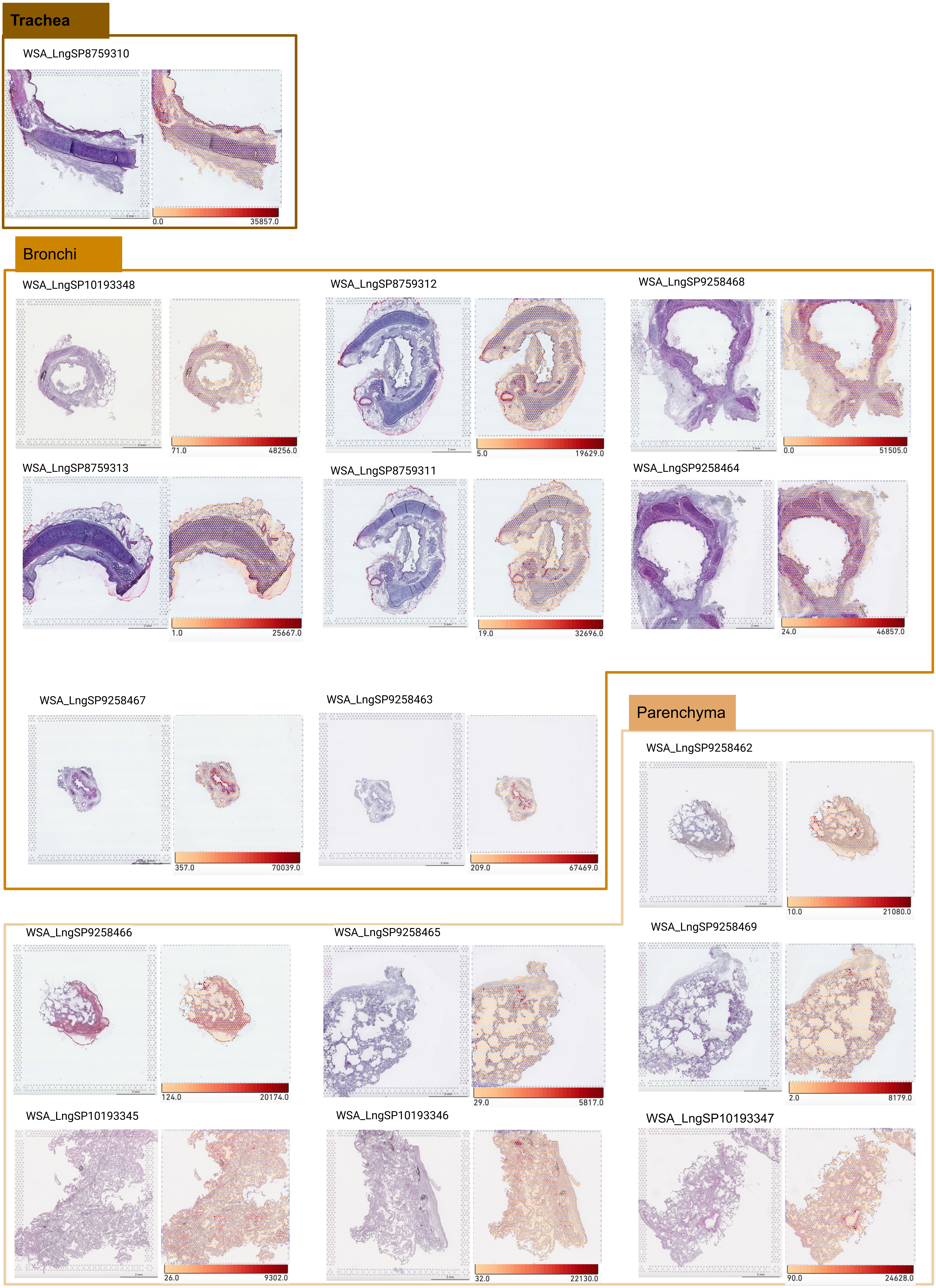

### Figure S3

**a**

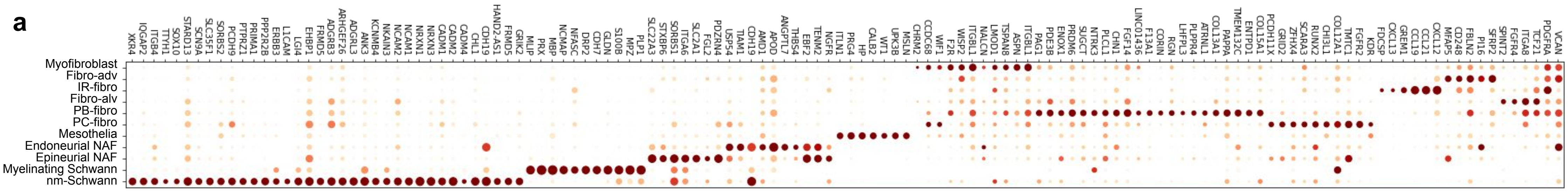


**b**

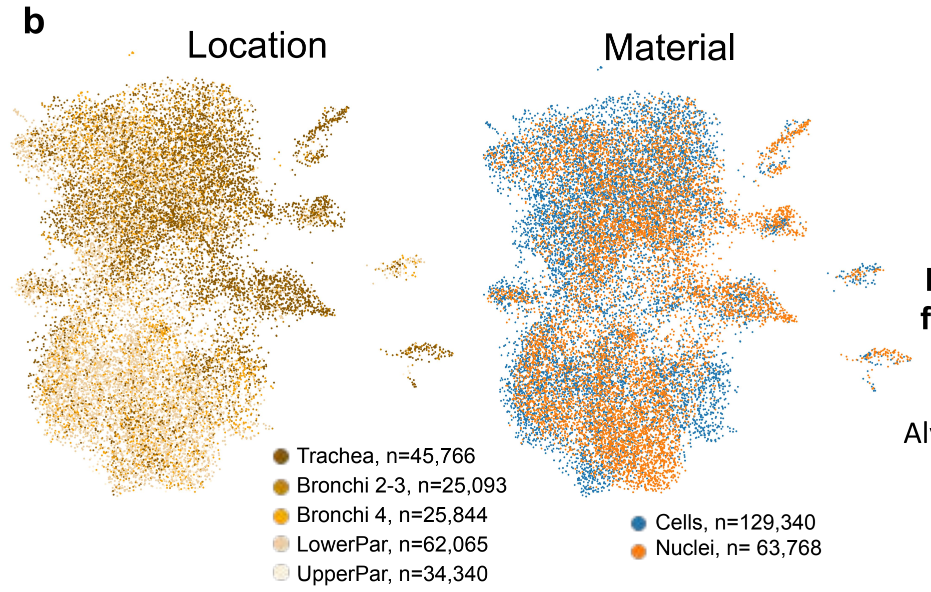


**c**

**Annotated cell type from current study**
**Total number of cell allocated to each cell type**

| Prediction cell type from Travaglini et al by Azimuth | Fibro-adventitial | Immune recruiting (IR) fibro | Fibro-alveolar | Myofibroblast | Perichondrial (PC) fibro | Peribronchial (PB) fibro | Mesothelial | Endoneurial NAF | Epineurial NAF | Myelinating Schwann | Nm Schwann |
|---|---|---|---|---|---|---|---|---|---|---|---|
| Alveolar Epithelial Type 2 | 0 | 0 | 1 | 0 | 0 | 0 | 0 | 0 | 0 | 0 | 0 |
| Bronchial Vessel | 1 | 0 | 0 | 0 | 0 | 0 | 0 | 0 | 0 | 0 | 0 |
| Basal | 4 | 0 | 1 | 0 | 0 | 0 | 0 | 0 | 0 | 1 | 0 |
| Smooth Muscle | 25 | 6 | 150 | 14 | 0 | 0 | 2 | 0 | 0 | 0 | 0 |
| Fibroblast | 6635 | 41 | 6333 | 160 | 345 | 11 | 24 | 25 | 19 | 0 | 3 |
| CD4 T | 232 | 0 | 52 | 11 | 15 | 40 | 6 | 0 | 38 | 7 | 30 |
| Capillary | 0 | 0 | 1 | 0 | 0 | 0 | 0 | 0 | 0 | 0 | 0 |
| Macrophage | 7 | 0 | 0 | 0 | 0 | 0 | 3 | 0 | 0 | 0 | 0 |
| Pericyte | 1644 | 3 | 488 | 67 | 196 | 288 | 7 | 41 | 89 | 0 | 76 |
| Myofibroblast | 539 | 1 | 379 | 590 | 511 | 68 | 15 | 3 | 4 | 0 | 0 |
| Lipofibroblast | 258 | 4 | 4 | 20 | 10 | 0 | 0 | 0 | 0 | 0 | 0 |
| Mesothelial | 825 | 4 | 11 | 8 | 6 | 4 | 83 | 5 | 15 | 0 | 0 |
| Neuroendocrine | 0 | 0 | 0 | 0 | 0 | 0 | 0 | 0 | 0 | 3 | 3 |

min ave max

Color - proportion of annotated cell type
Label - Number of cells in each category

**d**

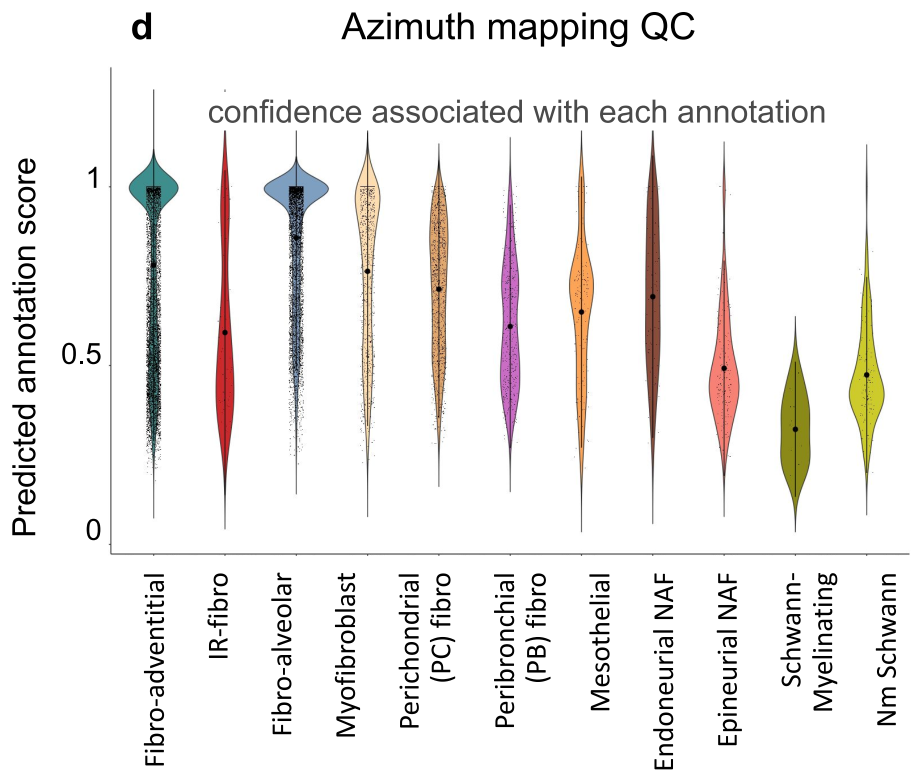

Figure S4

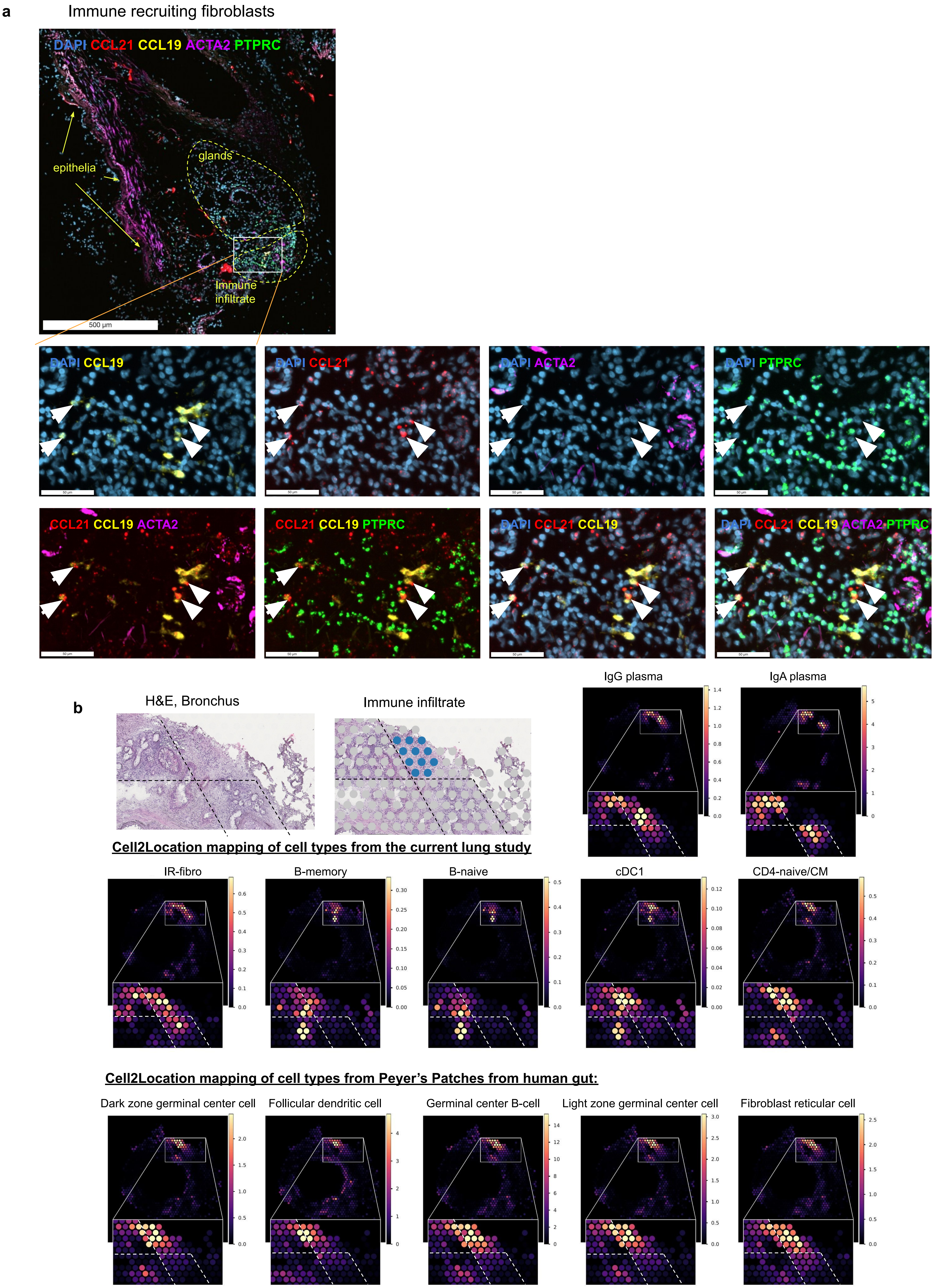

a
Immune recruiting fibroblasts
DAPI CCL21 CCL19 ACTA2 PTPRC
epithelia
glands
Immune infiltrate
500 µm
DAPI CCL19
DAPI CCL21
DAPI ACTA2
DAPI PTPRC
CCL21 CCL19 ACTA2
CCL21 CCL19 PTPRC
DAPI CCL21 CCL19
DAPI CCL21 CCL19 ACTA2 PTPRC
b
H&E, Bronchus
Immune infiltrate
IgG plasma
IgA plasma
Cell2Location mapping of cell types from the current lung study
IR-fibro
B-memory
B-naive
cDC1
CD4-naive/CM
Cell2Location mapping of cell types from Peyer's Patches from human gut:
Dark zone germinal center cell
Follicular dendritic cell
Germinal center B-cell
Light zone germinal center cell
Fibroblast reticular cell

Figure S5

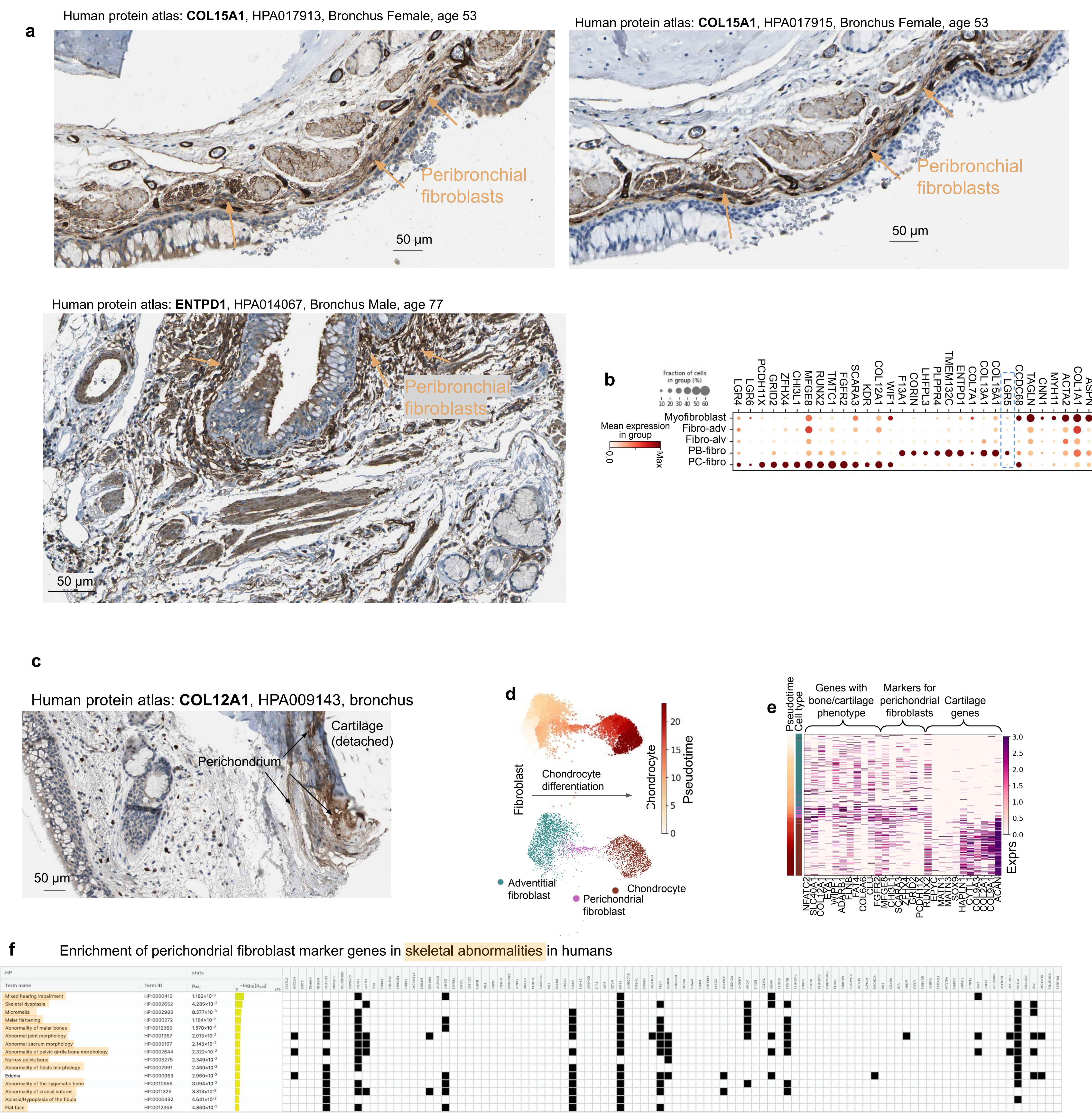
a
Human protein atlas: COL15A1, HPA017913, Bronchus Female, age 53
Peribronchial fibroblasts
50 µm
Human protein atlas: COL15A1, HPA017915, Bronchus Female, age 53
Peribronchial fibroblasts
50 µm
Human protein atlas: ENTPD1, HPA014067, Bronchus Male, age 77
Peribronchial fibroblasts
50 µm
b
Fraction of cells in group (%)
10 20 30 40 50 60
Mean expression in group
0.0
Max
Myofibroblast
Fibro-adv
Fibro-alv
PB-fibro
PC-fibro
LGR4
LGR6
PCDH11X
GRID2
ZFHX4
CHI3L1
MFGE8
RUNX2
TMTC1
FGFR2
SCARA3
KDR
COL12A1
WIF1
F13A1
CORIN
LHFPL3
PLPPR4
TMEM132C
ENTPD1
COL7A1
COL13A1
COL15A1
LGR5
CCDC68
TAGLN
CNN1
MYH11
ACTA2
COL1A1
ASPN
c
Human protein atlas: COL12A1, HPA009143, bronchus
Cartilage (detached)
Perichondrium
50 µm
d
Fibroblast
Chondrocyte differentiation
Chondrocyte
Pseudotime
0
5
10
15
20
Adventitial fibroblast
Perichondrial fibroblast
Chondrocyte
e
Pseudotime
Cell type
Genes with bone/cartilage phenotype
Markers for perichondrial fibroblasts
Cartilage genes
Exprs
3.0
2.5
2.0
1.5
1.0
0.5
0.0
NFATC2
SLC40A1
COL12A1
ELN
WIPF1
ADARB1
FLNB
FAT4
CLU
COL6A6
FGFR2
MFGE8
CHI3L1
SCARA3
ZFHX4
GRID2
PCDH11X
RUNX2
EYA1
MATN1
MATN3
SOX9
HAPLN1
COL9A3
COL2A1
COL9A1
ACAN
f
Enrichment of perichondrial fibroblast marker genes in skeletal abnormalities in humans
HP
stats
Term name
Term ID
$p_{adj}$
$-\log_{10}(p_{adj})$
Mixed hearing impairment
HP:0000410
$1.182\times10^{-3}$
Skeletal dysplasia
HP:0002652
$4.285\times10^{-3}$
Micromelia
HP:0002983
$8.577\times10^{-3}$
Malar flattening
HP:0000272
$1.184\times10^{-2}$
Abnormality of malar bones
HP:0012369
$1.570\times10^{-2}$
Abnormal joint morphology
HP:0001367
$2.015\times10^{-2}$
Abnormal sacrum morphology
HP:0005107
$2.145\times10^{-2}$
Abnormality of pelvic girdle bone morphology
HP:0002644
$2.322\times10^{-2}$
Narrow pelvis bone
HP:0003275
$2.349\times10^{-2}$
Abnormality of fibula morphology
HP:0002991
$2.460\times10^{-2}$
Edema
HP:0000969
$2.960\times10^{-2}$
Abnormality of the zygomatic bone
HP:0010668
$3.094\times10^{-2}$
Abnormality of cranial sutures
HP:0011329
$3.313\times10^{-2}$
Aplasia/Hypoplasia of the fibula
HP:0006492
$4.641\times10^{-2}$
Flat face
HP:0012368
$4.860\times10^{-2}$

Figure S6

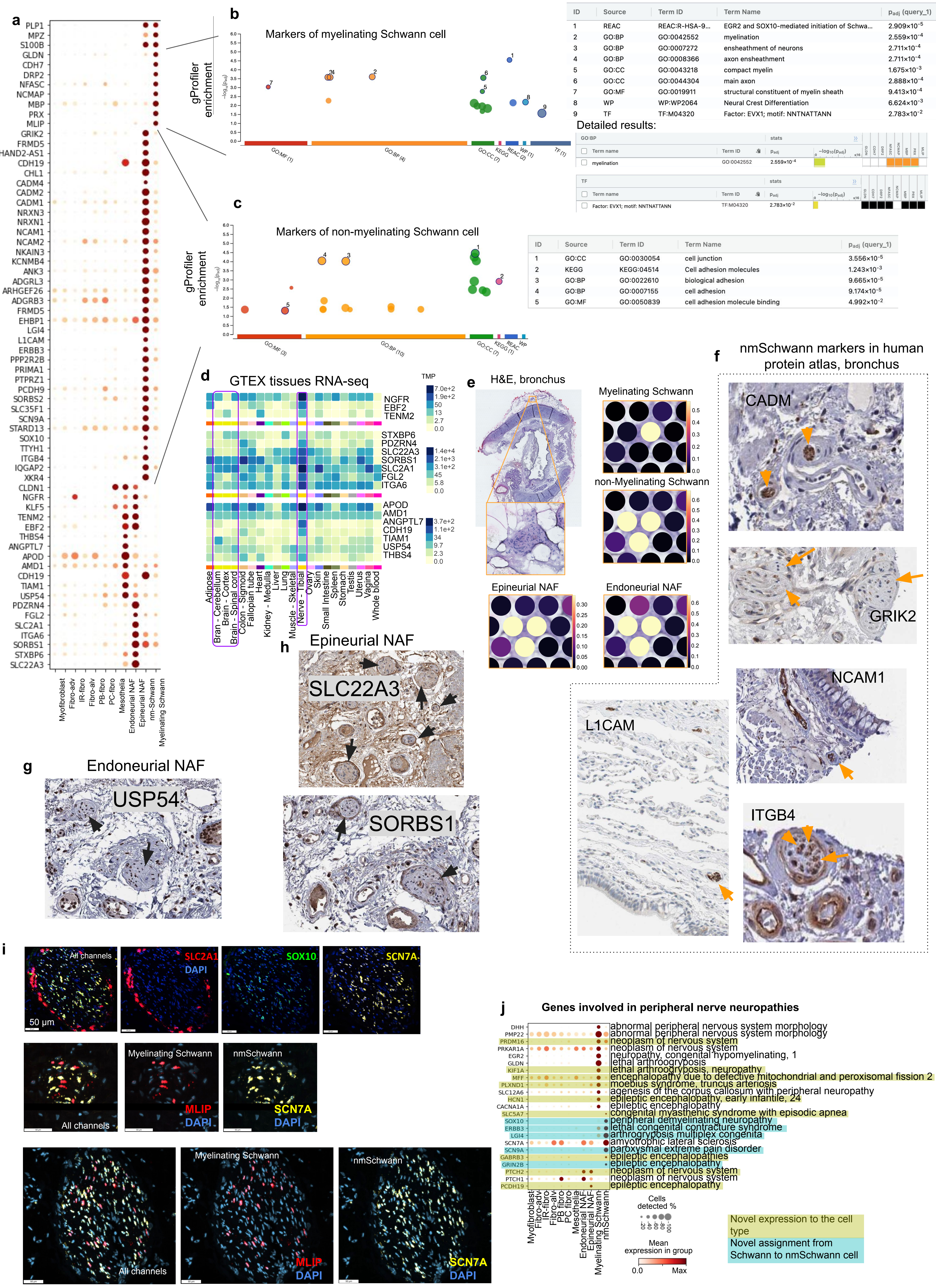
a
PLP1
MPZ
S100B
GLDN
CDH7
DRP2
NFASC
NCMAP
MBP
PRX
MLIP
GRIK2
FRMD5
HAND2-AS1
CDH19
CHL1
CADM4
CADM2
CADM1
NRXN3
NRXN1
NCAM1
NCAM2
NKAIN3
KCNMB4
ANK3
ADGRL3
ARHGEF26
ADGRB3
FRMD5
EHBP1
LGI4
L1CAM
ERBB3
PPP2R2B
PRIMA1
PTPRZ1
PCDH9
SORBS2
SLC35F1
SCN9A
STARD13
SOX10
TTYH1
ITGB4
IQGAP2
XKR4
CLDN1
NGFR
KLF5
TENM2
EBF2
THBS4
ANGPTL7
APOD
AMD1
CDH19
TIAM1
USP54
PDZRN4
FGL2
SLC2A1
ITGA6
SORBS1
STXBP6
SLC22A3
Myofibroblast
Fibro-adv
IR-fibro
Fibro-alv
PB-fibro
PC-fibro
Mesothelia
Endoneurial NAF
Epineurial NAF
nm-Schwann
Myelinating Schwann
b
Markers of myelinating Schwann cell
gProfiler enrichment
GO:MF (1)
GO:BP (4)
GO:CC (2)
KEGG
REAC (2)
WP (1)
TF (1)
ID | Source | Term ID | Term Name | $p_{adj}$ (query_1)
1 | REAC | REAC:R-HSA-9... | EGR2 and SOX10-mediated initiation of Schwa... | $2.909\times10^{-5}$
2 | GO:BP | GO:0042552 | myelination | $2.559\times10^{-4}$
3 | GO:BP | GO:0007272 | ensheathment of neurons | $2.711\times10^{-4}$
4 | GO:BP | GO:0008366 | axon ensheathment | $2.711\times10^{-4}$
5 | GO:CC | GO:0043218 | compact myelin | $1.675\times10^{-3}$
6 | GO:CC | GO:0044304 | main axon | $2.888\times10^{-4}$
7 | GO:MF | GO:0019911 | structural constituent of myelin sheath | $9.413\times10^{-4}$
8 | WP | WP:WP2064 | Neural Crest Differentiation | $6.624\times10^{-3}$
9 | TF | TF:M04320 | Factor: EVX1; motif: NNTNATTANN | $2.783\times10^{-2}$
Detailed results:
GO:BP
Term name | Term ID | $p_{adj}$ | $-\log_{10}(p_{adj})$
myelination | GO:0042552 | $2.559\times10^{-4}$
TF
Term name | Term ID | $p_{adj}$ | $-\log_{10}(p_{adj})$
Factor: EVX1; motif: NNTNATTANN | TF:M04320 | $2.783\times10^{-2}$
c
Markers of non-myelinating Schwann cell
gProfiler enrichment
GO:MF (3)
GO:BP (10)
GO:CC (7)
KEGG (1)
REAC
WP
ID | Source | Term ID | Term Name | $p_{adj}$ (query_1)
1 | GO:CC | GO:0030054 | cell junction | $3.556\times10^{-5}$
2 | KEGG | KEGG:04514 | Cell adhesion molecules | $1.243\times10^{-3}$
3 | GO:BP | GO:0022610 | biological adhesion | $9.665\times10^{-5}$
4 | GO:BP | GO:0007155 | cell adhesion | $9.174\times10^{-5}$
5 | GO:MF | GO:0050839 | cell adhesion molecule binding | $4.992\times10^{-2}$
d
GTEX tissues RNA-seq
TMP
NGFR
EBF2
TENM2
STXBP6
PDZRN4
SLC22A3
SORBS1
SLC2A1
FGL2
ITGA6
APOD
AMD1
ANGPTL7
CDH19
TIAM1
USP54
THBS4
Adipose
Bran - Cerebellum
Brain - Cortex
Brain - Spinal cord
Colon - Sigmoid
Fallopian tube
Heart
Kidney - Medulla
Liver
Lung
Muscle - Skeletal
Nerve - Tibial
Ovary
Skin
Small Intestine
Spleen
Stomach
Testis
Uterus
Vagina
Whole blood
e
H&E, bronchus
Myelinating Schwann
non-Myelinating Schwann
Epineurial NAF
Endoneurial NAF
f
nmSchwann markers in human protein atlas, bronchus
CADM
GRIK2
NCAM1
L1CAM
ITGB4
g
Endoneurial NAF
USP54
h
Epineurial NAF
SLC22A3
SORBS1
i
All channels
SLC2A1
DAPI
SOX10
SCN7A
50 µm
Myelinating Schwann
nmSchwann
MLIP
j
Genes involved in peripheral nerve neuropathies
DHH
PMP22
PRDM16
PRKAR1A
EGR2
GLDN
KIF1A
MFF
PLXND1
SLC12A6
HCN1
CACNA1A
SLC5A7
SOX10
ERBB3
LGI4
SCN7A
SCN9A
GABRB3
GRIN2B
PTCH2
PTCH1
PCDH19
abnormal peripheral nervous system morphology
abnormal peripheral nervous system morphology
neoplasm of nervous system
neoplasm of nervous system
neuropathy, congenital hypomyelinating, 1
lethal arthrogryposis
lethal arthrogryposis, neuropathy
encephalopathy due to defective mitochondrial and peroxisomal fission 2
moebius syndrome, truncus arteriosis
agenesis of the corpus callosum with peripheral neuropathy
epileptic encephalopathy, early infantile, 24
epileptic encephalopathy
congenital myasthenic syndrome with episodic apnea
peripheral demyelinating neuropathy
lethal congenital contracture syndrome
arthrogryposis multiplex congenita
amyotrophic lateral sclerosis
paroxysmal extreme pain disorder
epileptic encephalopathies
epileptic encephalopathy
neoplasm of nervous system
neoplasm of nervous system
epileptic encephalopathy
Myofibroblast
Fibro-adv
IR-fibro
Fibro-alv
PB fibro
PC fibro
Mesothelia
Endoneurial NAF
Epineurial NAF
Myelinating Schwann
nmSchwann
Cells detected %
Mean expression in group
0.0
Max
Novel expression to the cell type
Novel assignment from Schwann to nmSchwann cell

Figure S7

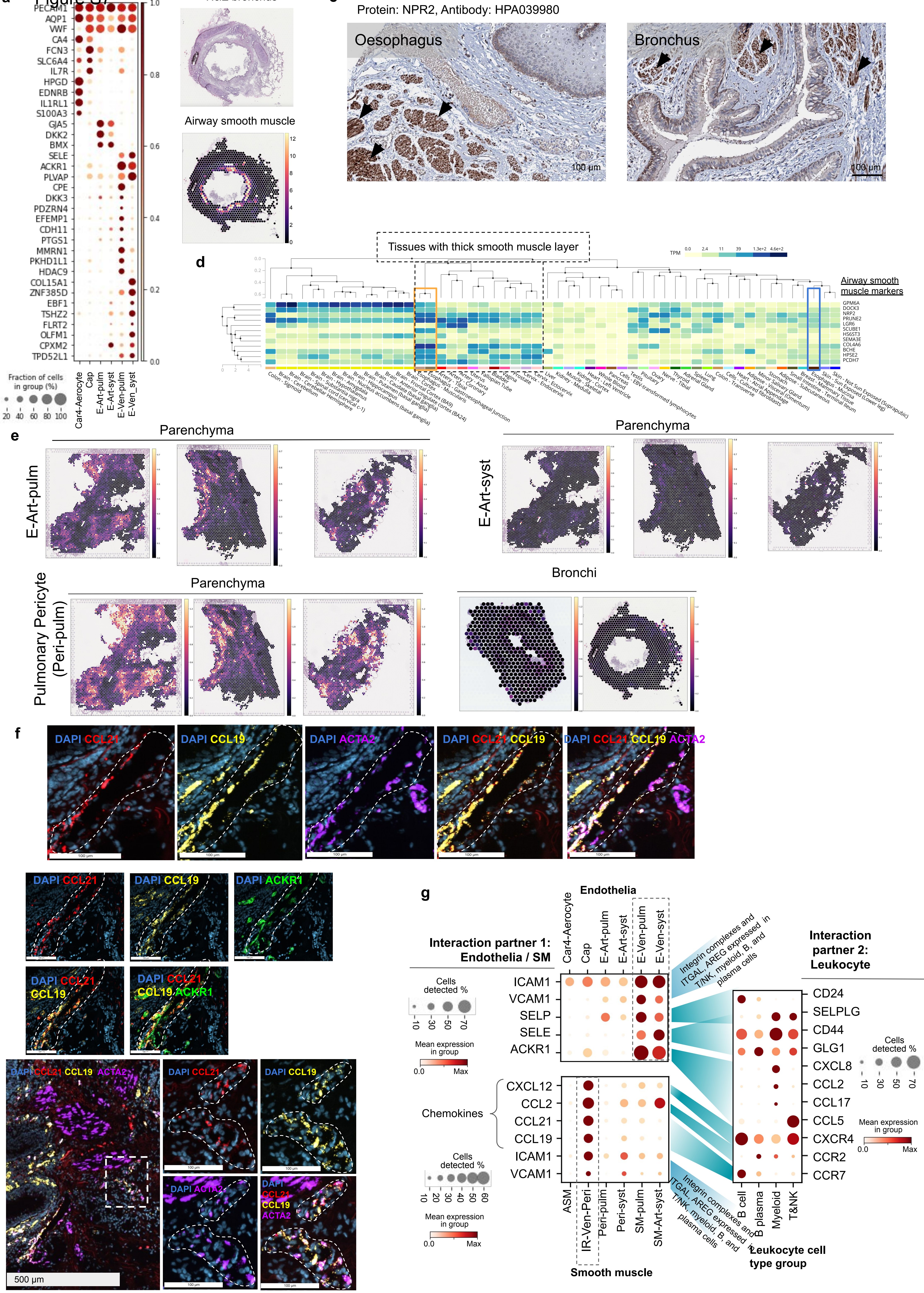
a
PECAM1
AQP1
VWF
CA4
FCN3
SLC6A4
IL7R
HPGD
EDNRB
IL1RL1
S100A3
GJA5
DKK2
BMX
SELE
ACKR1
PLVAP
CPE
DKK3
PDZRN4
EFEMP1
CDH11
PTGS1
MMRN1
PKHD1L1
HDAC9
COL15A1
ZNF385D
EBF1
TSHZ2
FLRT2
OLFM1
CPXM2
TPD52L1
Fraction of cells in group (%)
20 40 60 80 100
Car4-Aerocyte
Cap
E-Art-pulm
E-Art-syst
E-Ven-pulm
E-Ven_syst
1.0 0.8 0.6 0.4 0.2 0.0
b
H&E bronchus
Airway smooth muscle
c
Protein: NPR2, Antibody: HPA039980
Oesophagus
Bronchus
100 µm
100 µm
d
Tissues with thick smooth muscle layer
TPM
Airway smooth muscle markers
GPM6A
DOCK3
NPR2
PRUNE2
LGR6
SCUBE1
HS6ST3
SEMA3E
COL4A6
BCHE
HPSE2
PCDH7
e
Parenchyma
E-Art-pulm
Parenchyma
E-Art-syst
Parenchyma
Pulmonary Pericyte (Peri-pulm)
Bronchi
f
DAPI CCL21
DAPI CCL19
DAPI ACTA2
DAPI CCL21 CCL19
DAPI CCL21 CCL19 ACTA2
DAPI CCL21
DAPI CCL19
DAPI ACKR1
DAPI CCL21 CCL19
DAPI CCL21 CCL19 ACKR1
DAPI CCL21 CCL19 ACTA2
DAPI CCL21
DAPI CCL19
DAPI ACTA2
DAPI CCL21 CCL19 ACTA2
500 µm
g
Interaction partner 1: Endothelia / SM
Endothelia
Car4-Aerocyte
Cap
E-Art-pulm
E-Art-syst
E-Ven-pulm
E-Ven-syst
Cells detected %
10 30 50 70
Mean expression in group
0.0 Max
ICAM1
VCAM1
SELP
SELE
ACKR1
Chemokines
CXCL12
CCL2
CCL21
CCL19
ICAM1
VCAM1
Cells detected %
10 20 30 40 50 60
Mean expression in group
0.0 Max
ASM
IR-Ven-Peri
Peri-pulm
Peri-syst
SM-pulm
SM-Art-syst
Smooth muscle
Integrin complexes and ITGAL, AREG expressed in T/NK, myeloid, B, and plasma cells
Integrin complexes and ITGAL, AREG expressed in T/NK, myeloid, B, and plasma cells
Interaction partner 2: Leukocyte
CD24
SELPLG
CD44
GLG1
CXCL8
CCL2
CCL17
CCL5
CXCR4
CCR2
CCR7
B cell
B plasma
Myeloid
T&NK
Leukocyte cell type group
Cells detected %
10 30 50 70
Mean expression in group
0.0 Max

### Figure S8

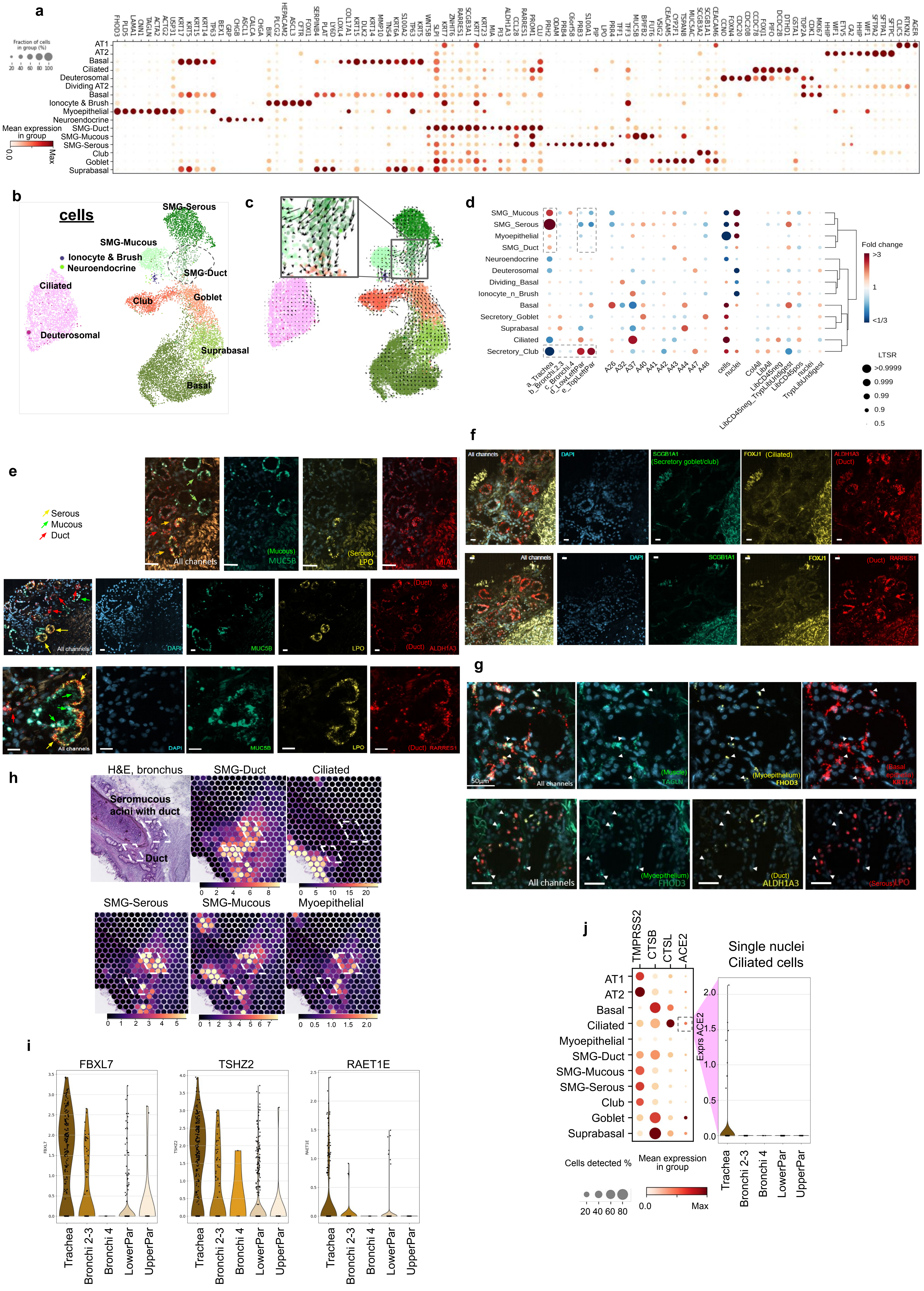
a
Fraction of cells in group (%)
Mean expression in group
AT1
AT2
Basal
Ciliated
Deuterosomal
Dividing AT2
Ionocyte & Brush
Myoepithelial
Neuroendocrine
SMG-Duct
SMG-Mucous
SMG-Serous
Club
Goblet
Suprabasal
b
cells
SMG-Serous
SMG-Mucous
Ionocyte & Brush
Neuroendocrine
SMG-Duct
Ciliated
Club
Goblet
Deuterosomal
Suprabasal
Basal
c
d
Fold change
LTSR
e
Serous
Mucous
Duct
All channels
(Mucous) MUC5B
(Serous) LPO
MIA
DAPI
(Duct) ALDH1A3
(Duct) RARRES1
f
All channels
DAPI
SCGB1A1 (Secretory goblet/club)
FOXJ1 (Ciliated)
ALDH1A3 (Duct)
(Duct) RARRES1
g
50μm
(Muscle) TAGLN
(Myoepithelium) FHOD3
(Basal epithelia) KRT14
(Duct) ALDH1A3
(Serous) LPO
h
H&E, bronchus
Seromucous acini with duct
Duct
SMG-Duct
Ciliated
SMG-Serous
SMG-Mucous
Myoepithelial
i
FBXL7
TSHZ2
RAET1E
Trachea
Bronchi 2-3
Bronchi 4
LowerPar
UpperPar
j
TMPRSS2
CTSB
CTSL
ACE2
Single nuclei Ciliated cells
Exprs ACE2
Cells detected %
Mean expression in group
Max

### Figure S9

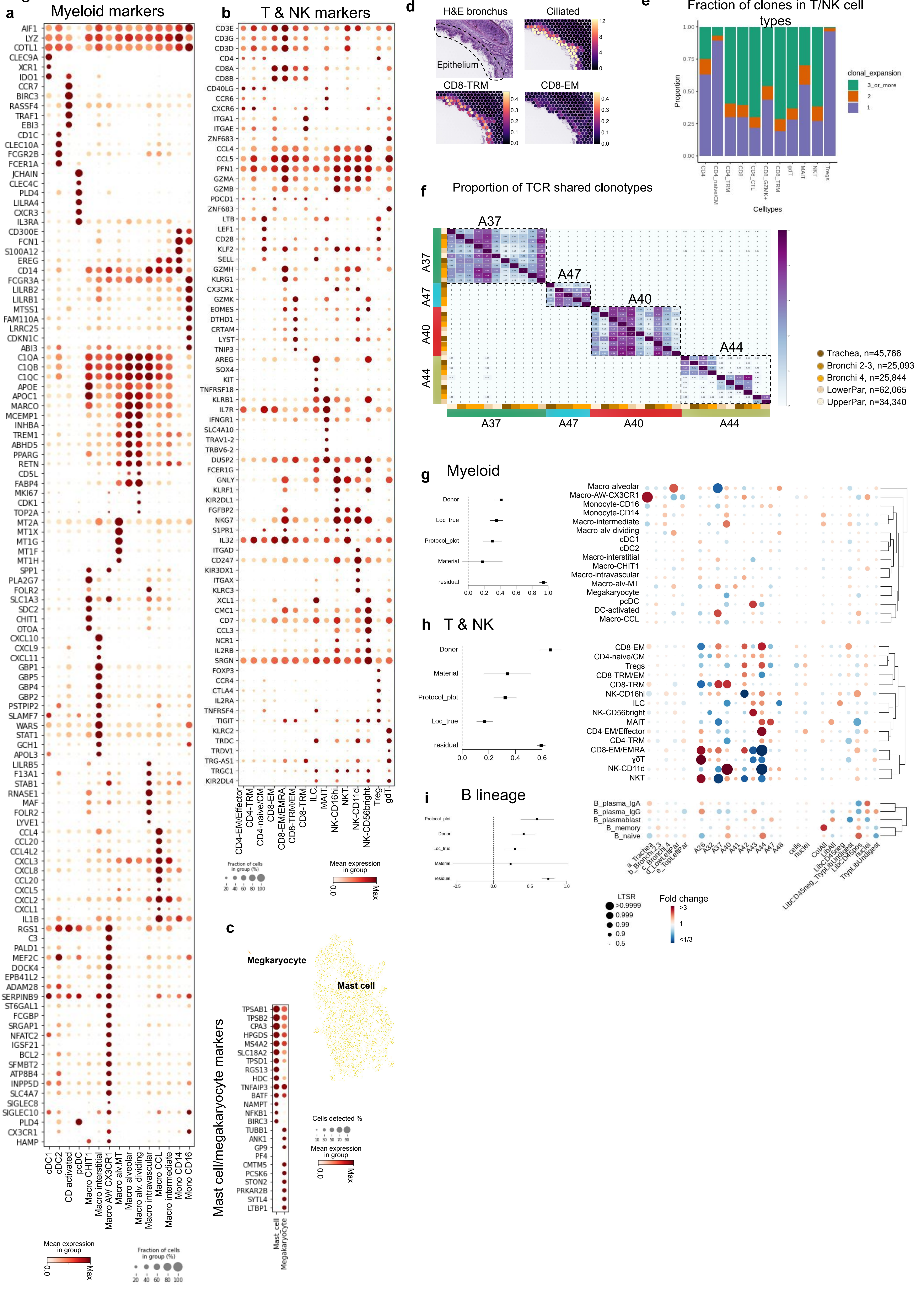
a Myeloid markers
b T & NK markers
c Mast cell/megakaryocyte markers
Megakaryocyte
Mast cell
d H&E bronchus
Ciliated
Epithelium
CD8-TRM
CD8-EM
e Fraction of clones in T/NK cell types
clonal_expansion
3_or_more
2
1
Proportion
Celltypes
f Proportion of TCR shared clonotypes
A37
A47
A40
A44
Trachea, n=45,766
Bronchi 2-3, n=25,093
Bronchi 4, n=25,844
LowerPar, n=62,065
UpperPar, n=34,340
g Myeloid
h T & NK
i B lineage
Donor
Loc_true
Protocol_plot
Material
residual
LTSR
Fold change
Mean expression in group
Fraction of cells in group (%)
Cells detected %

Figure S10

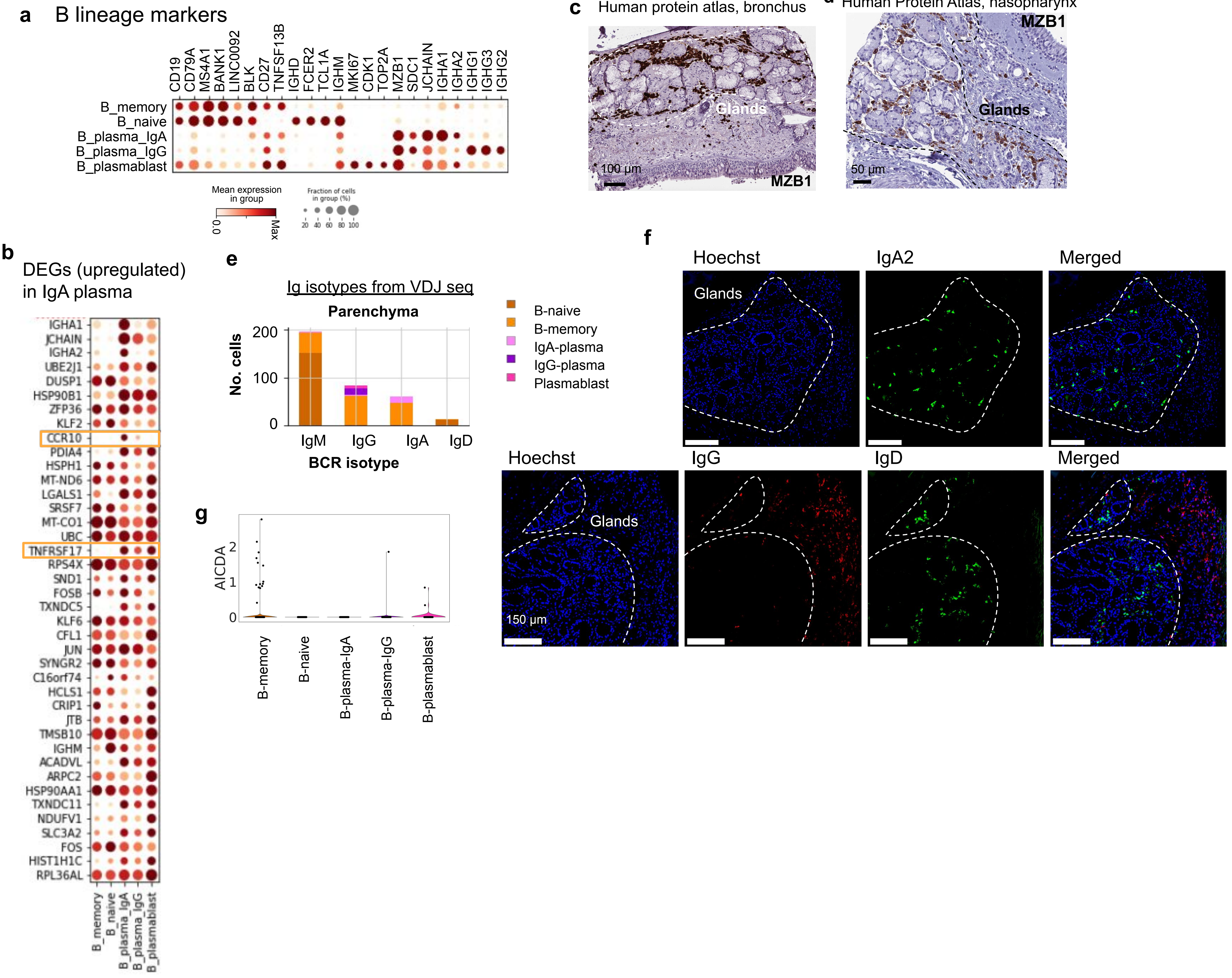
a
B lineage markers
CD19
CD79A
MS4A1
BANK1
LINC0092
BLK
CD27
TNFSF13B
IGHD
FCER2
TCL1A
IGHM
MKI67
CDK1
TOP2A
MZB1
SDC1
JCHAIN
IGHA1
IGHA2
IGHG1
IGHG3
IGHG2
B_memory
B_naive
B_plasma_IgA
B_plasma_IgG
B_plasmablast
Mean expression in group
0.0
Max
Fraction of cells in group (%)
20 40 60 80 100
c
Human protein atlas, bronchus
Glands
100 µm
MZB1
d
Human Protein Atlas, nasopharynx
MZB1
Glands
50 µm
b
DEGs (upregulated) in IgA plasma
IGHA1
JCHAIN
IGHA2
UBE2J1
DUSP1
HSP90B1
ZFP36
KLF2
CCR10
PDIA4
HSPH1
MT-ND6
LGALS1
SRSF7
MT-CO1
UBC
TNFRSF17
RPS4X
SND1
FOSB
TXNDC5
KLF6
CFL1
JUN
SYNGR2
C16orf74
HCLS1
CRIP1
JTB
TMSB10
IGHM
ACADVL
ARPC2
HSP90AA1
TXNDC11
NDUFV1
SLC3A2
FOS
HIST1H1C
RPL36AL
B_memory
B_naive
B_plasma_IgA
B_plasma_IgG
B_plasmablast
e
Ig isotypes from VDJ seq
Parenchyma
No. cells
200
100
0
IgM
IgG
IgA
IgD
BCR isotype
B-naive
B-memory
IgA-plasma
IgG-plasma
Plasmablast
f
Hoechst
IgA2
Merged
Glands
Hoechst
IgG
IgD
Merged
Glands
150 µm
g
AICDA
2
1
0
B-memory
B-naive
B-plasma-IgA
B-plasma-IgG
B-plasmablast
